# SpatioCAD: Context-aware graph diffusion model for pinpointing spatially variable genes in heterogeneous tissues

**DOI:** 10.64898/2026.03.06.708087

**Authors:** Hao Wen, Qunlun Shen, Shuqin Zhang

## Abstract

Spatial transcriptomics enables comprehensive characterization of tissue architecture, and the identification of spatially variable genes (SVGs) is a critical step for defining region-specific molecular markers and uncovering spatially regulated mechanisms across diverse biological contexts. However, most existing methods for SVG detection overlook cell density variations, a major confounding factor in complex tissues such as tumors, where heterogeneous cellularity frequently introduces false-positive calls. Here we present SpatioCAD, a computational framework that explicitly decouples genuine spatial expression patterns from confounding effects driven by cellularity. SpatioCAD leverages and extends a graph diffusion model to simulate expression propagation under cell-density-aware conditions, thereby ensuring unbiased detection of SVGs across all expression levels. Systematic evaluations on simulated datasets demonstrate its superior statistical power and specificity. Applied to breast cancer, lung cancer, and glioma datasets, SpatioCAD identifies functionally diverse SVGs, including low-abundance transcripts with established roles in tumor progression, while also recapitulates biologically meaningful tissue architecture features.

## 1 Introduction

Spatial transcriptomics (ST) technologies enable the comprehensive gene expression profiling at near- or sub-cellular resolution within the native tissue context, providing unprecedented insights into the spatial organization of cells, their in situ functional states, the cell-cell communications and spatially distinct gene expression programs [1–4]. They have been widely applied to unravel the complex cellular interactions and delineate spatially organized functional domains that govern a broad spectrum of fundamental biological processes, from development to disease [5–8]. In particular, ST facilitates the systematic characterization of pathological tissue architectures, such as tumor microenvironments and spatially restricted malignant niches, through the analysis of spatially resolved human disease atlases [8].

A fundamental task in spatial transcriptomics data analysis is the identification of genes exhibiting spatially structured expression patterns [4, 9]. These genes, known as spatially variable genes (SVGs), play a critical role in deciphering functionally diverse tissue regions and revealing the key molecular drivers of biological processes [10, 11]. For instance, SVG analysis has uncovered the distinct spatial-pathological mechanisms underlying amyloid-*β* and *τ* proteins in Alzheimer’s disease [12]. Consequently, numerous computational methods have been proposed for SVG identification, each founded on distinct mathematical and statistical principles. For example, SPARK-X [13] and SpatialDE [4] employ statistical frameworks, such as Gaussian process regression and nonparametric approaches, to test spatial variance and dependence of gene expression; SpaGFT [14] utilizes a graph Fourier transform on spatial neighbor networks to decompose gene expression signals into frequency components, effectively highlighting regionally structured variation; Sepal [15] leverages diffusion-based smoothing and spatial autocorrelation analysis to enhance the detection of genes exhibiting coherent local expression domains. Additional scalable approaches such as nnSVG [16] and SOMDE [17] have further extended SVG identification to increasingly complex and large-scale datasets.

While existing SVG identification methods have been proven effective in well-structured tissues, their reliability is severely compromised in highly heterogeneous contexts such as the tumor microenvironment (TME). A key feature of the TME is the dramatic spatial variation in cell density resulting from the uncontrolled proliferation of cancer cells [18], which can be misinterpreted as genuine biological signal, thereby constituting a major confounding factor for SVG identification. Failing to account for this confounder risks missing key drivers of tissue function. Although one recent method, STMiner, employs a framework based on Optimal Transport (OT) to address this challenge [19], its high computational cost and sensitivity to outliers limit broader application [20, 21]. Furthermore, a prevalent bias in many current algorithms is the tendency to favor genes with high expression levels, frequently leading to the exclusion of key regulatory genes that are often expressed at low levels [22].

To overcome these challenges, we present SpatioCAD (SPAtially variable gene identification using Context-Aware graph Diffusion model), a computational framework built on our proposed Node-Attributed Graph Diffusion (NAGD) model to explicitly decouple genuine spatial patterns from cellular density variations. When benchmarked on simulated datasets, SpatioCAD achieves superior statistical power and effective false discovery rate control across diverse spatial configurations and expression patterns. Applied to human breast and lung cancer datasets, SpatioCAD demonstrates exceptional accuracy, unbiasedness and spatial coherence through multidimensional evaluations. Biologically, SpatioCAD identifies SVGs strongly associated with tumor aggressiveness, immune infiltration, and patient prognosis. Furthermore, when applied to a diffuse midline glioma (DMG) sample, SpatioCAD enables the precise delineation of key histological regions, including the tumor core, invasive margin, and peritumoral reactive zone. These results collectively establish SpatioCAD as a powerful and robust tool for deciphering the spatial architecture of complex tissues and offering novel insights into their underlying biology.

## 2 Results

### 2.1 SpatioCAD overview

SpatioCAD is a computational framework designed for the unbiased and efficient identification of SVGs within highly heterogeneous tissues, such as the tumor microenvironment. A critical confounder in these contexts is spatial variation in cell density, which, if not properly addressed, frequently leads to false-positive SVG calls. To overcome this challenge, SpatioCAD is built on two fundamental assumptions: first, biologically meaningful spatial patterns manifest as locally smooth signals; and second, structured signals require a longer diffusion time to reach a steady state than random ones.

Leveraging the first assumption, SpatioCAD first employs a noise filtering module to exclude genes dominated by high-frequency variation. Specifically, we propose a metric called Roughness Score to quantify the magnitude of signal variation during the initial phase of a standard graph diffusion process (Fig. 1A). The theoretical foundation of this metric is that genes with structured spatial patterns exhibit minimal signal variations during early diffusion due to their local consistency, while noise genes change drastically. This distinction allows SpatioCAD to adaptively filter out uninformative genes, thereby preventing them from confounding downstream analysis.

**Figure 1.**
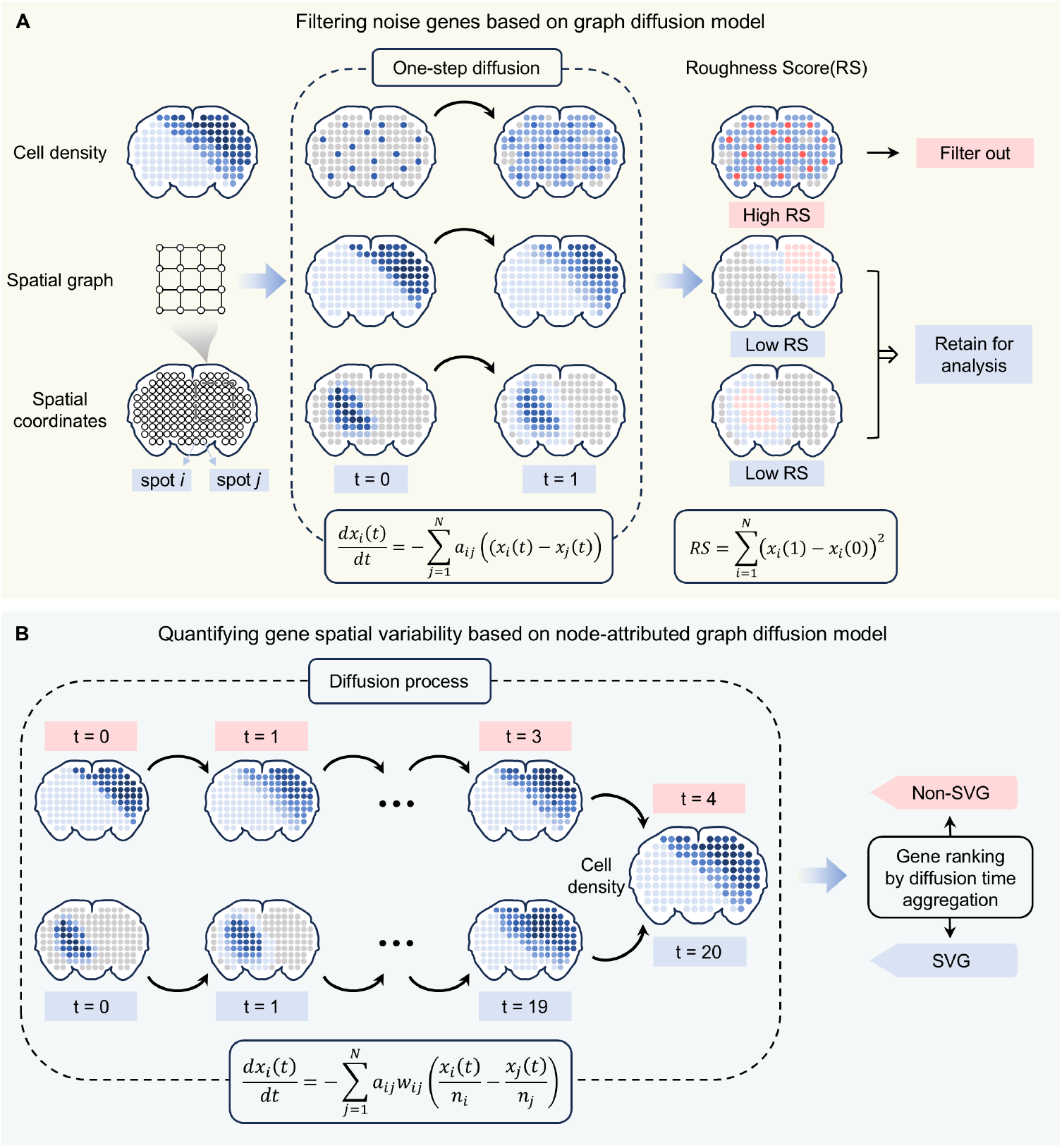
Workflow of SpatioCAD. SpatioCAD is composed of two main modules: noise gene filtering and SVG identification. **A** SpatioCAD first constructs a spatial graph representing the spatial relationships between locations and calculates the cell density for each location using the input spatial transcriptomics data. Then, SpatioCAD filters out noise genes based on the difference between the initial gene expression profile and the profile after a one-step diffusion. **B** SpatioCAD identifies SVGs by computing the diffusion time to the steady state (proportional to cell density) under the Node-Attributed Graph Diffusion (NAGD) model. Genes requiring a longer time to reach the steady state are considered to exhibit significant spatial variation.

Next, SpatioCAD leverages our proposed Node-Attributed Graph Diffusion (NAGD) model to rank the spatial variability of remaining genes based on their characteristic diffusion times (Fig. 1B). NAGD generalizes the standard graph diffusion by guiding the diffusion process along per-cell signal gradients rather than relying on absolute signal differences between spots, thereby decoupling genuine spatial patterns from cellular density variations. Building on the second assumption, we then quantify the spatial variability of each gene by measuring the time required to reach a steady state. A key theoretical foundation of SpatioCAD is that all normalized signals within the NAGD framework converge to a steady state proportional to cell density (Supplementary Note). This convergence property ensures that the diffusion time serves as a universally comparable metric across genes, enabling a robust and unbiased ranking of SVGs.

### 2.2 Simulation analysis

To systematically benchmark the performance of SpatioCAD against existing methods, we generated datasets simulating the cellular heterogeneity characteristic of tumor tissues (**Methods**). We designed two different domain settings of tissue slices (Fig. 2A, first row). Each slice contained five distinct domains, with domain E designated as the tumor core characterized by high cell density. The gene expression counts were generated by sampling from a Zero-Inflated Negative Binomial (ZINB) distribution with different fold change parameters, which were modeled as a product of cell density and spatial variability. The cell density varied between domains, and was higher in the tumor core (Fig. 2A, second row). The spatial variability for background genes was uniform across domains, whereas for SVGs, it was set to be higher or lower within the specific domains. The resulting final fold change thus captured their compounded effect (Fig. 2B). For each simulated tissue architecture, we generated 900 background genes and 200 SVGs, and considered two cases with and without 50 noise genes. We note that although background genes may appear to exhibit spatial patterns, these patterns are driven by cellular heterogeneity and are conceptually distinct from true SVGs (Fig. 2C, Supplementary Fig. 1).

**Figure 2.**
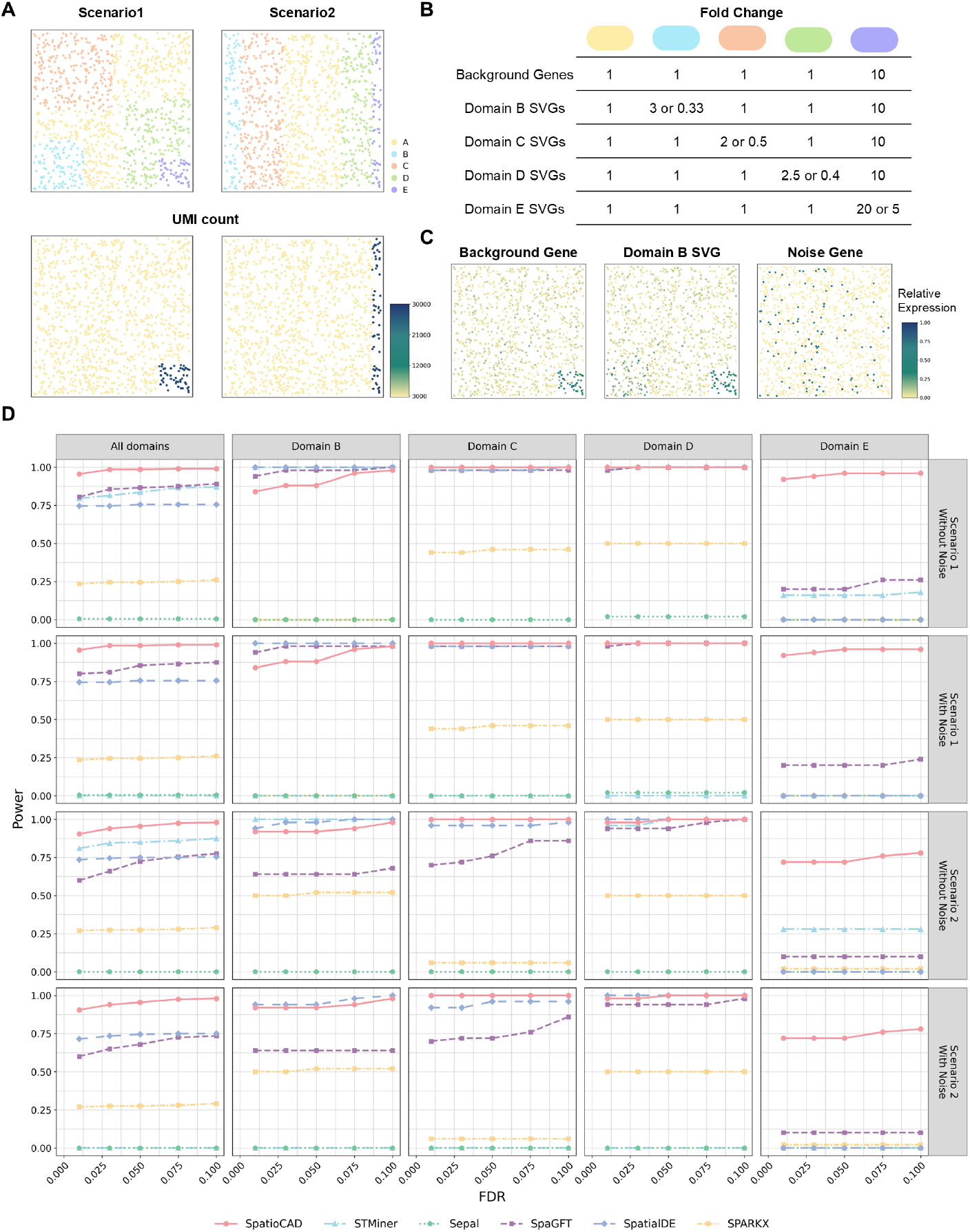
Simulation framework and benchmarking results. **A** Top: two simulated tissue layouts. Scenario 1 with square-shaped domains and Scenario 2 with strip-shaped domains. Colors represent distinct spatial domains. Bottom: spatial distribution of UMI counts corresponding to the two scenarios. **B** Simulation parameter settings of fold-change for different gene categories (**Methods**). **C** Representative spatial expression patterns in Scenario 1 for a background gene, a domain-specific SVG, and a noise gene. **D** Power plots show the statistical power (y-axis) detected by different methods (represented by different colors and line types) at a range of FDR (x-axis) across all domains and the four separate domains (columns) and the two scenarios with and without noise (rows).

We benchmarked SpatioCAD against STMiner [19], Sepal [15], SpaGFT [14], SpatialDE [4], and SPARK-X [13]. To account for the diverse output statistics generated by these methods, we evaluated performance by calculating their statistical power at a range of predefined False Discovery Rates (FDRs) to ensure a fair comparison (**Methods**).

We first assessed the global performance of each method on simulated datasets without noise genes (Fig. 2D, left column). SpatioCAD consistently surpassed all other competing methods, achieving the highest statistical power. While STMiner and SpatialDE demonstrated stability across the two simulation settings, SpaGFT exhibited notable volatility: it performed competitively in Scenario 1 but declined substantially in Scenario 2. SPARK-X showed limited detection capability, while Sepal displayed negligible sensitivity. Specifically, at an FDR threshold of 0.1, SpatioCAD achieved a statistical power of 0.99 (Scenario 1) and 0.98 (Scenario 2), substantially surpassing the second-to fourth-best methods: STMiner (0.87 and 0.88), SpaGFT (0.88 and 0.74), and SpatialDE (0.76 and 0.75). By contrast, the remaining methods yielded powers below 0.30 (SPARK-X: 0.26 and 0.29; Sepal: 0.005 and 0), indicating their limited efficacy under these challenging conditions. The excellent efficacy of SpatioCAD and STMiner is attributable to their specialized designs for complex tissues, as both account for cellular density heterogeneity. And SpatioCAD was more powerful than STMiner by explicitly modeling the diffusion process at the individual cell level, rather than at the location level, thereby achieving finer signal resolution. In comparison, for SpaGFT and SpatialDE, though they normalized cellular expression levels at each location to mitigate density effects, they failed to capture the intrinsic impact of varying cell counts. The performance fluctuations in SpaGFT may stem from the spectral characteristics of graph Fourier transform-based methods, where the capture efficiency for specific spatial patterns depends on their alignment with the graph Laplacian’s eigenvectors. An analysis of SPARK-X revealed that it identified nearly all background genes and SVGs as spatially significant (*p <* 0.05, Supplementary Fig. 2). This suggests that SPARK-X misinterprets density-driven heterogeneity as true biological signal. Similarly, an examination of Sepal’s gene ranking showed a complete conflation of background genes and SVGs (Supplementary Fig. 3). This failure likely results from Sepal’s diffusion model assuming a uniform steady state, which fails to account for the confounding effect of cellular density.

We next evaluated the robustness of each method by introducing noise genes into the simulation. While most methods maintained stable performance, STMiner showed a significant performance decline, with its statistical power dropping from approximately 0.87 in noise-free conditions to nearly zero at an FDR threshold of 0.1 upon noise inclusion. Further investigation of the results revealed that STMiner tends to falsely rank noise genes as top candidates (Supplementary Fig. 4), corroborating the known sensitivity of Optimal Transport-based methods to expression outliers [20, 21]. Although stringent quality control in real-world datasets might mitigate this issue, the consistently high performance of SpatioCAD underscores its inherent robustness and reduced reliance on ideal preprocessing.

We further examined the performance of each method across distinct spatial domains. Given the minimal impact of noise on most methods’ rankings and the limited global performance of SPARK-X and Sepal, we focused our detailed domain-level analysis on the remaining methods under noise-free conditions. Domains B, C, and D represent normal tissue regions with relatively uniform cell density, making SVG detection analogous to standard spatial analysis. In Domains C and D, SpatioCAD, STMiner, and SpatialDE all demonstrated robust performance across two scenarios, achieving near-perfect power (approximately 1.0) across all FDR thresholds (Fig. 2D, third and fourth columns from the left). In Domain B, while STMiner and SpatialDE maintained perfect performance, SpatioCAD exhibited a marginal decline. This slight difference is intrinsic to the method’s design: it models diffusion at the individual cell level, rather than smoothing expression over locations. This may modestly reduce the sensitivity in regions with low total cell counts, such as Domain B. However, its maintained power of 0.98 at an FDR threshold of 0.1 confirms its exceptional reliability under these constraints. Additionally, SpaGFT continued to show scenario-dependent fluctuations across these domains, likely due to the variable alignment between spatial patterns and the graph Laplacian’s eigenvectors.

The most significant performance divergence occurred in Domain E (tumor core), where high cell density creates a strong confounding background. Accurately detecting SVGs in this region is particularly challenging, requiring the capacity to decouple genuine spatial expression from density-driven variation. SpatioCAD was the only method that succeeded in this task (Fig. 2D, right column), achieving a power of 0.96 (Scenario 1) and 0.78 (Scenario 2) at an FDR threshold of 0.1, whereas all other methods attained powers below 0.3. This highlights a critical limitation in competing methods: while they may partially adjust for cell density, they fail to disentangle the compounded effects when spatial variability and density variation co-occur. In contrast, SpatioCAD’s superior performance stems from its NAGD framework, which by directly modeling diffusion at the cellular level, inherently accounts for uneven cell densities and thereby avoids the signal distortion inherent in conventional location-level normalization.

In summary, the simulation analyses not only confirm SpatioCAD as a powerful and robust SVG detection method but also offer a critical benchmarking framework to address the pervasive challenge of cellular heterogeneity in spatial transcriptomics.

### 2.3 Evaluation of SpatioCAD on human breast cancer data reveals unbiased and biologically informative detection of SVGs

We applied SpatioCAD along with the five previously mentioned methods to analyze a public 10X Visium dataset of human breast cancer [23], which comprised 18,085 genes measured across 4,992 spots. Accounting for the diverse output statistics across methods, we focused the comparative analysis on the top 1,000 SVGs identified by each approach to ensure a standardized and equitable assessment. Within the tissue, cancer-associated regions, including invasive regions and DCIS, exhibited higher UMI counts (Fig. 3A), aligning well with the established biological prior of increased cell density within tumor regions [24], and highlighting cellularity as a critical confounding factor in spatial transcriptomic analysis.

**Figure 3.**
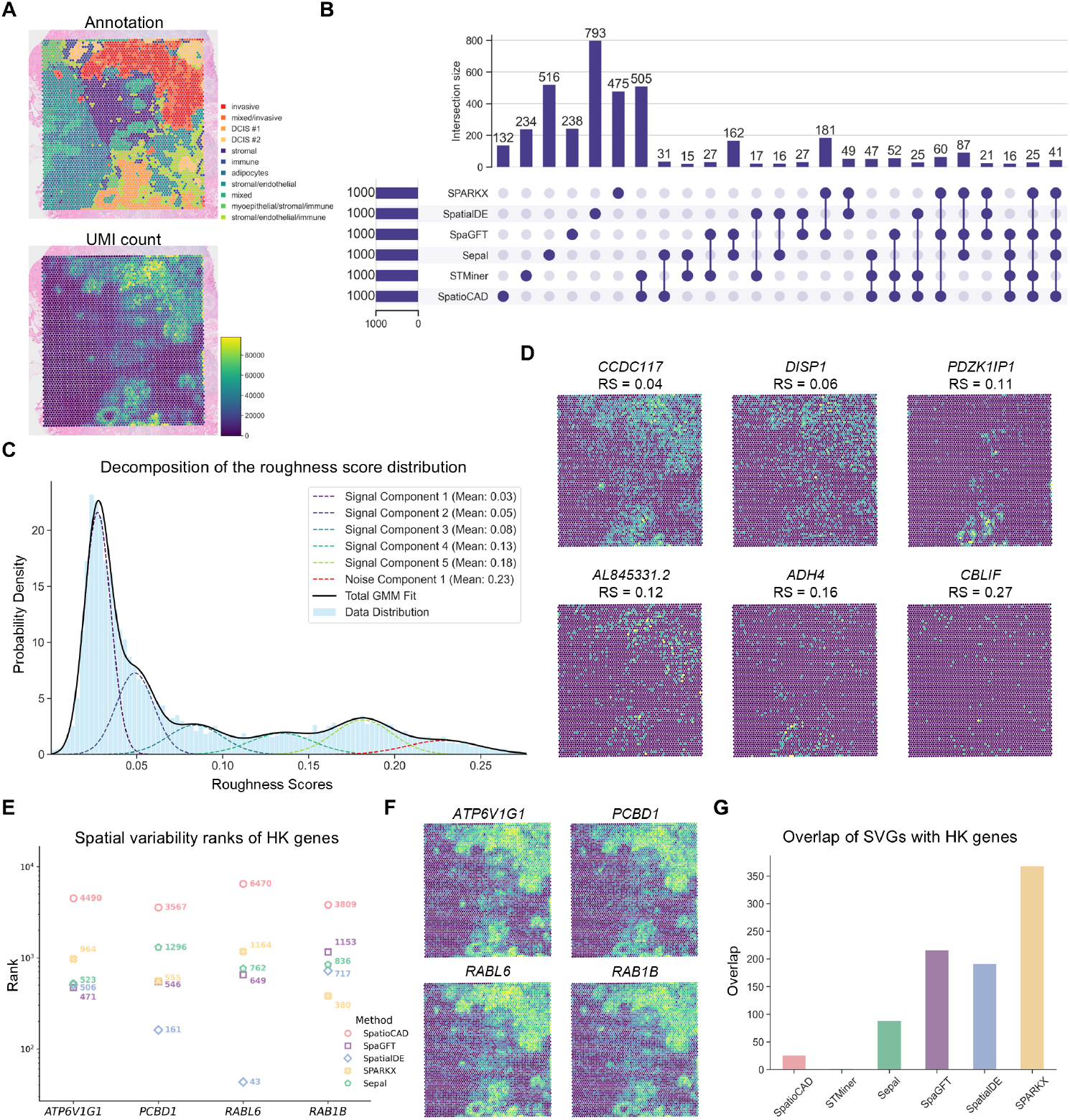
SpatioCAD decouples biological spatial variability from cell density effects in human breast cancer data. **A** Visualization of the human breast cancer dataset with its annotations (upper) and UMI counts (lower). **B** UpSet plot shows the overlap among the top 1,000 SVGs identified by the six compared methods (SpatioCAD, STMiner, Sepal, SpaGFT, SpatialDE, SPARK-X). **C** Decomposition of the Roughness Score (RS) distribution for all genes using a Gaussian Mixture Model, identifying six distinct components in the human breast cancer data. **D** Spatial expression patterns of representative genes from each component with their corresponding RSs. **E** Comparison of spatial variability ranks for representative housekeeping genes when using different methods. **F** Spatial expression patterns of representative house-keeping genes shown in **E. G** The number of overlaps between the top 1,000 SVGs identified by different methods and known housekeeping genes in public databases.

#### 2.3.1 SpatioCAD effectively decouples biological spatial variability from tissue heterogeneity

To comprehensively compare the results of the six methods, we generated an UpSet plot depicting the overlaps among their respective top SVGs (Fig. 3B), which illustrates the distinct gene preferences of different methods. Notably, SpatioCAD and STMiner shared the largest intersection (505 genes), attributable to their capacity to adjust for cellular heterogeneity. Other pronounced overlaps were also observed, such as between SpaGFT and SPARK-X (181 genes), and between SpaGFT and Sepal (162 genes). Furthermore, SpatioCAD identified the fewest unique SVGs (132 genes), suggesting high detection reliability as most SVGs were corroborated by other methods.

We employed Roughness Score (RS) developed from standard graph diffusion model to filter out noise genes (**Methods**). All genes were partitioned into six components according to RS, with the component possessing the highest mean RS being designated as the noise (Fig. 3C). This classification was supported by visualization: genes from the highest-RS component (e.g., *CBLIF*) displayed disordered distributions, while genes from lower-RS components (e.g., *ADH4, CCDC117*) showed structured patterns (Fig. 3D). These results confirm the efficacy of the RS as a metric for noise gene filtering, thereby enhancing the statistical power for subsequent SVG identification.

We evaluated the quality of the identified SVGs by using housekeeping (HK) genes as negative controls [25], as they are expected to lack spatial variability. Examination of several HK genes with markedly different rankings between methods revealed that many were spuriously identified as top SVGs by conventional approaches, whereas SpatioCAD correctly excluded them by ranking them beyond the top 1000 (Fig. 3E). As shown in Fig. 3F, the spatial expression of these genes strongly correlated with UMI counts, confirming their density-dependent nature. Quantitatively, the overlap between the SVGs and HK genes was lowest for STMiner (2 genes) and comparably low for SpatioCAD (26 genes). Both methods significantly outperformed the others, which showed substantially higher overlaps: Sepal (88 genes), SpaGFT (216 genes), SpatialDE (191 genes), and SPARK-X (368 genes) (Fig. 3G). This finding emphasizes the importance of accounting for cell density for capturing genuine spatial variation in heterogeneous tissues.

#### 2.3.2 SpatioCAD eliminates expression-level bias and uncovers functionally diverse SVGs

Given the substantial divergence in SVG lists, we investigated whether the top SVGs identified by each method were systematically skewed toward higher or lower expression by calculating their normalized expression ranks relative to all genes (**Methods**). Ideally, an unbiased method should yield SVGs with normalized expression ranks centered around 0.5. Quantitative analysis revealed that SpatioCAD achieved the least bias (Fig. 4A), with a mean normalized rank of 0.487, and a mean absolute deviation (MAD) of 0.013 from the ideal value of 0.5. In contrast, STMiner showed a preference for lowly expressed genes (mean: 0.398; MAD: 0.102), whereas SpatialDE (mean: 0.687; MAD: 0.187), Sepal (mean: 0.859; MAD: 0.359), SpaGFT (mean: 0.871; MAD: 0.371), and SPARK-X (mean: 0.904; MAD: 0.404) were heavily biased towards highly expressed genes. Spearman correlation analysis between spatial variability scores and expression levels further supported these findings (**Methods**): SpatioCAD displayed negligible correlation (*R* = −0.04), while STMiner displayed a significant negative correlation (*R* = −0.58). All others displayed significant positive correlations, ranging from SpatialDE (*R* = 0.71) to SpaGFT (*R* = 0.83), SPARK-X (*R* = 0.90), and Sepal (*R* = 0.92). Visual inspection of rank scatter plots corroborated these metrics: SVGs identified by SpatioCAD were uniformly distributed across expression levels, while the SVGs from STMiner clustered along anti-diagonal and those from the remaining methods clustered along diagonal patterns indicating systematic bias toward either low or high expression (Fig. 4B, Supplementary Fig. 5). These results confirm that SpatioCAD effectively mitigates expression-level confounding, ensuring unbiased SVG identification.

**Figure 4.**
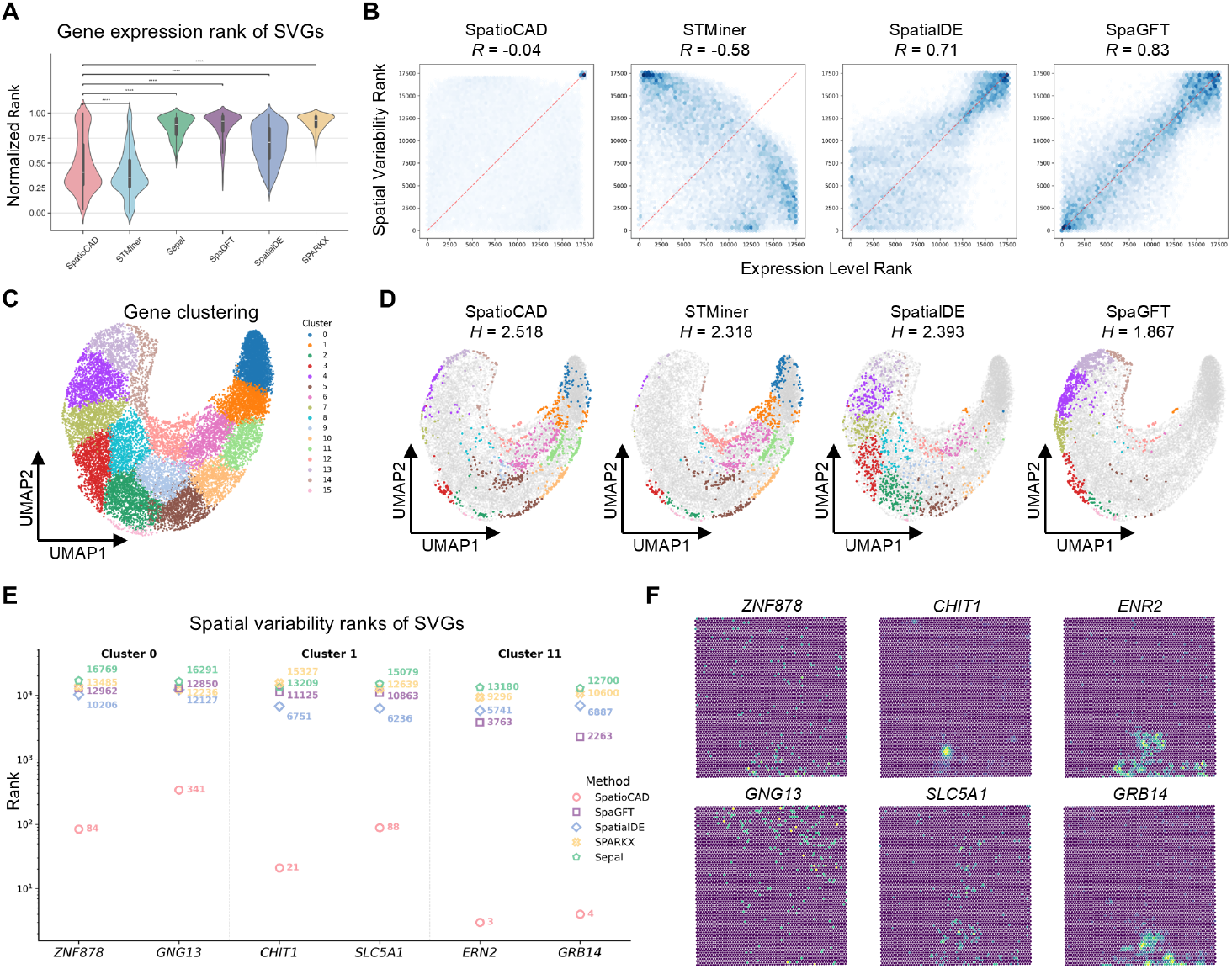
SpatioCAD identifies functionally diverse SVGs with minimal expression bias in human breast cancer data. **A** Violin plot shows the distribution of normalized expression ranks (y-axis) for the top 1,000 SVGs identified by each method. Wilcoxon rank-sum tests were applied to normalized expression ranks comparing SpatioCAD with other methods (*p*-values = 3.37 × 10^−10^, 3.30 × 10^−162^, 3.66 × 10^−170^, 2.03 × 10^−67^, 6.94 × 10^−190^, respectively). **B** Hexbin density plots show the relationship between gene expression ranks (x-axis) and spatial variability ranks (y-axis). Spearman correlation between the two ranks was calculated for each method (SpatioCAD: −0.04, STMiner: −0.58, SpatialDE: 0.71, SpaGFT: 0.83, SPARK-X: 0.90, Sepal: 0.92). **C** UMAP plot shows the clustering results of all genes based on their expression profiles. **D** Projection of the top 1,000 SVGs identified by each method onto the global gene UMAP. Shannon entropy of their distribution across these clusters was calculated for each method (SpatioCAD: 2.518, STMiner: 2.318, SpatialDE: 2.393, SpaGFT: 1.867, SPARK-X: 1.449, Sepal: 2.013) with higher value indicating more functionally diverse SVGs. **E** Comparison of spatial variability ranks for representative genes from clusters 0, 1, and 11. SpatioCAD ranks these genes higher than other methods (SpatialDE, SpaGFT, SPARK-X, Sepal). **F** Spatial expression patterns of representative genes shown in **E**.

To illustrate the advantages of unbiased detection, we evaluated the functional diversity of the SVGs identified by each method. All genes were first grouped into distinct clusters based on their expression profiles (Fig. 4C). Gene set enrichment analysis using the Gene Ontology (GO) database confirmed the biological relevance of each cluster (Supplementary Figs. 6-8), and the cluster size and expression levels were quantified (Supplementary Figs. 9-10). We then calculated the Shannon entropy of the SVG distribution across these clusters (**Methods**). A method that captures functionally diverse SVGs should distribute them broadly across clusters, resulting in higher entropy. SpatioCAD achieved the highest entropy (*H* = 2.518), followed by SpatialDE (*H* = 2.393), STMiner (*H* = 2.318), Sepal (*H* = 2.013), SpaGFT (*H* = 1.867), and SPARK-X (*H* = 1.449) (Fig. 4D, Supplementary Fig. 11). This indicates that SpatioCAD captures a significantly broader spectrum of biologically functional signals compared to other methods.

A detailed breakdown of SVG counts per cluster further confirmed the distinct biases for each method (Supplementary Fig. 12). While SpatioCAD successfully detected biological signals from each cluster, SpatialDE, Sepal, SpaGFT, and SPARK-X identified few or no SVGs in clusters 0, 1, and 11, which correspond to the lowest average gene expression levels (Supplementary Fig. 9). Conversely, STMiner showed limited sensitivity in clusters 4, 7, 8, 9, and 13, consistent with its bias toward lowly expressed genes. These observations align with the earlier correlation analysis between spatial variability and gene expression levels, underscoring the expression-level bias of these methods in detecting spatial patterns.

We further validated the biological relevance of several SVGs from low-expression clusters (Clusters 0, 1, and 11) that were identified by SpatioCAD but missed by other methods. Their relative rankings across methods are shown in Fig. 4E. In Cluster 0, *ZNF878* exhibited a spatial distribution pattern matching the DCIS subtype (Fig. 4F, Fig. 3A). This observation aligns with the biological context of zinc finger proteins, a family increasingly recognized for their potential roles in cancer progression [26]. *GNG13* exhibited distinct spatial clustering in the invasive regions (Fig. 4F). As a member of the G protein *γ* subunit family, it acts as a critical signal transducer for chemokine receptors [27]. This enrichment at the invasive regions likely reflects the active recruitment of migratory machinery in these disseminating tumor cells. In Cluster 1, *CHIT1* exhibited a highly focal expression pattern within the DCIS regions (Fig. 4F). Since *CHIT1* is a biochemical marker for activated macrophages [28], this local region likely reflects tumor-associated macrophage infiltration. Moreover, *SLC5A1* (*SGLT1*) was broadly enriched in DCIS and invasive regions (Fig. 4F), consistent with the high glucose uptake required for the glycolytic metabolism of rapidly proliferating pre-invasive cells [29]. In Cluster 11, *ERN2* and *GRB14* exhibited spatial patterns that closely mirrored the DCIS regions (Fig. 4F). This co-localization highlights the complex physiological state of pre-invasive tumor cells. Specifically, the high expression of *ERN2*, an endoplasmic reticulum (ER) stress sensor, reflects the activation of the unfolded protein response (UPR) triggered by the intense biosynthetic load and proteotoxic stress inherent to rapidly growing tumor nests [30]. In parallel, *GRB14* functions as a key signal modulator driving breast cancer cell cycle progression and proliferation [31]. These validations demonstrate that SpatioCAD can reliably identify functionally relevant spatial patterns even among low-abundance transcripts, uncovering biological signals that are otherwise overlooked by expression-biased methods.

#### 2.3.3 SpatioCAD captures more robust biological structures compared to STMiner

We performed a direct comparison between SpatioCAD and STMiner, a leading approach that addresses cellularity, and focused on the SVGs uniquely identified by each method. After removing common SVGs, both methods yielded an equal number of exclusive candidates. We first assessed spatial coherence of the SVGs using a local linear regression model that predicts gene expression at a specific location based on its neighboring coordinates, to quantify local smoothness, a characteristic of genuine biological signals (**Methods**). The SVGs identified by SpatioCAD exhibited a significantly lower prediction error compared to those from STMiner (MAE: 3.15 vs 4.95, Wilcoxon-rank test *p*-value = 8.00 × 10^−26^; Fig. 5A). This result indicates that SpatioCAD detects spatial patterns with high spatial continuity, effectively filtering out the spatially incoherent noise.

**Figure 5.**
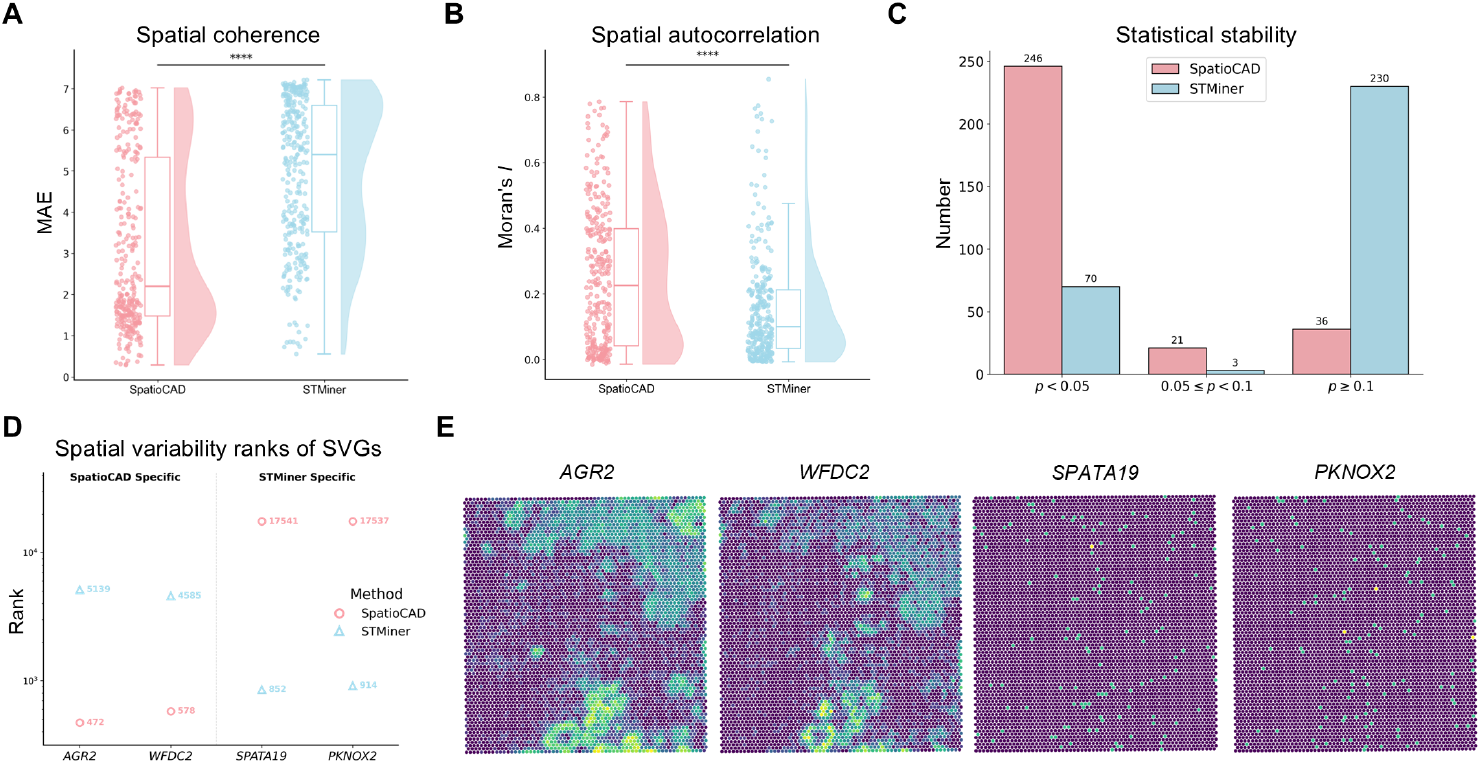
Comparison between SpatioCAD and STMiner for SVG identification in human breast cancer data. **A** Raincloud plots show the distribution of the gene expression reconstruction error (MAE) for method-specific SVGs. Wilcoxon rank-sum test comparing SpatioCAD with STMiner (*p*-value = 8.00 × 10^−26^). **B** Raincloud plot shows the distribution of global Moran’s *I* indices for method-specific SVGs. Wilcoxon rank-sum test comparing SpatioCAD with STMiner (*p*-value = 1.72 × 10^−7^). **C** Bar plot shows the classification of method-specific SVGs based on permutationbased empirical *p*-values. **D** Comparison of spatial variability ranks for representative method-specific SVGs. **E** Spatial expression patterns of representative genes shown in **D**.

We then evaluated the global spatial structure of the identified SVGs using Moran’s *I*, a standard measure of spatial autocorrelation [32]. A higher Moran’s *I* indicates stronger spatial clustering of expression, as opposed to a random or dispersed distribution. SVGs from SpatioCAD consistently displayed significantly higher Moran’s *I* indices (mean: 0.25 vs 0.15, Wilcoxon-rank test *p*-value = 1.72 × 10^−7^; Fig. 5B). This result confirms that SpatioCAD captures genes with structured spatial patterns, whereas STMiner may select genes with weak or spatially fragmented distributions.

We further assessed the statistical stability of SVG detection using permutation-based test (**Methods**) to distinguish true biological variability from random background fluctuations. SpatioCAD yielded significantly lower empirical *p*-values for its top-ranked SVGs compared to STMiner (Fig. 5C). Specifically, 246 SVGs identified by SpatioCAD were statistically significant (*p <* 0.05), whereas only 70 of STMiner’s top SVGs met this threshold. Conversely, a substantial number of genes identified by STMiner (230 genes) exhibited high *p*-values (*p* ≥ 0.1), suggesting they may be false positives driven by noise. This result suggests that the signals captured by SpatioCAD represent statistically robust biological spatial features rather than stochastic fluctuations.

To visually inspect the discrepancies between the two methods, we examined the spatial expression patterns of genes with significantly different ranks (Fig. 5D). SpatioCAD successfully prioritized biologically relevant markers, such as *AGR2* and *WFDC2. AGR2* is a well-established estrogen-regulated biomarker and potential therapeutic target in breast cancer [33], while *WFDC2* (*HE4*) has been associated with lymph node metastasis and poor disease-free survival in breast cancer patients [34]. Visual inspection also confirmed that these genes exhibit structured and continuous spatial patterns (Fig. 5E, left). In contrast, STMiner prioritized some genes characterized by poor spatial predictability and low statistical confidence, such as *SPATA19* (*p*-value = 0.90) and *PKNOX2* (*p*-value = 0.92), which displayed scattered, discrete noise patterns lacking spatial coherence (Fig. 5E, right). Functional analysis also implies that these identifications are likely false positives driven by technical noise. For instance, *SPATA19* is known to be associated with spermatogenesis [35], which is biologically irrelevant to breast tissue.

For the computational efficiency, as STMiner relies on computationally intensive Optimal Transport (OT) to model cellularity, SpatioCAD demonstrated a substantial improvement, completing the analysis in 43 seconds compared to 41,983 seconds for STMiner. This acceleration is directly attributable to the analytical solution of diffusion time in the NAGD model, which enables rapid computation without iterative approximations.

Collectively, these quantitative and qualitative assessments demonstrate SpatioCAD’s superior ability to distinguish biologically meaningful spatial signals from technical artifacts.

### 2.4 SpatioCAD demonstrates unbiased and biologically informative detection of SVGs in human lung cancer data

To assess the generalizability of our findings across different tissue architectures, we applied SpatioCAD and the competing methods to analyze a public 10X Visium dataset of human lung cancer, which comprised 18,085 genes measured across 3,858 spots (Fig. 6A). To ensure an equitable comparison, we also focused our analysis on the top 1,000 SVGs identified by each method. The observed heterogeneity in UMI counts across spots further reflects the marked variation in cellular density within this tissue.

#### 2.4.1 SpatioCAD effectively corrects for tissue heterogeneity, minimizes expression-level bias and recovers functionally diverse SVGs

We generated an UpSet plot depicting the overlaps among the respective top SVGs identified by the compared methods (Fig. 6B). The largest overlap was observed between SpatioCAD and STMiner (531 genes), attributable to their shared capability to adjust for cellular heterogeneity. SpatioCAD identified the fewest unique SVGs (156 genes), indicating the high reliability as most of its findings were supported by other methods.

**Figure 6.**
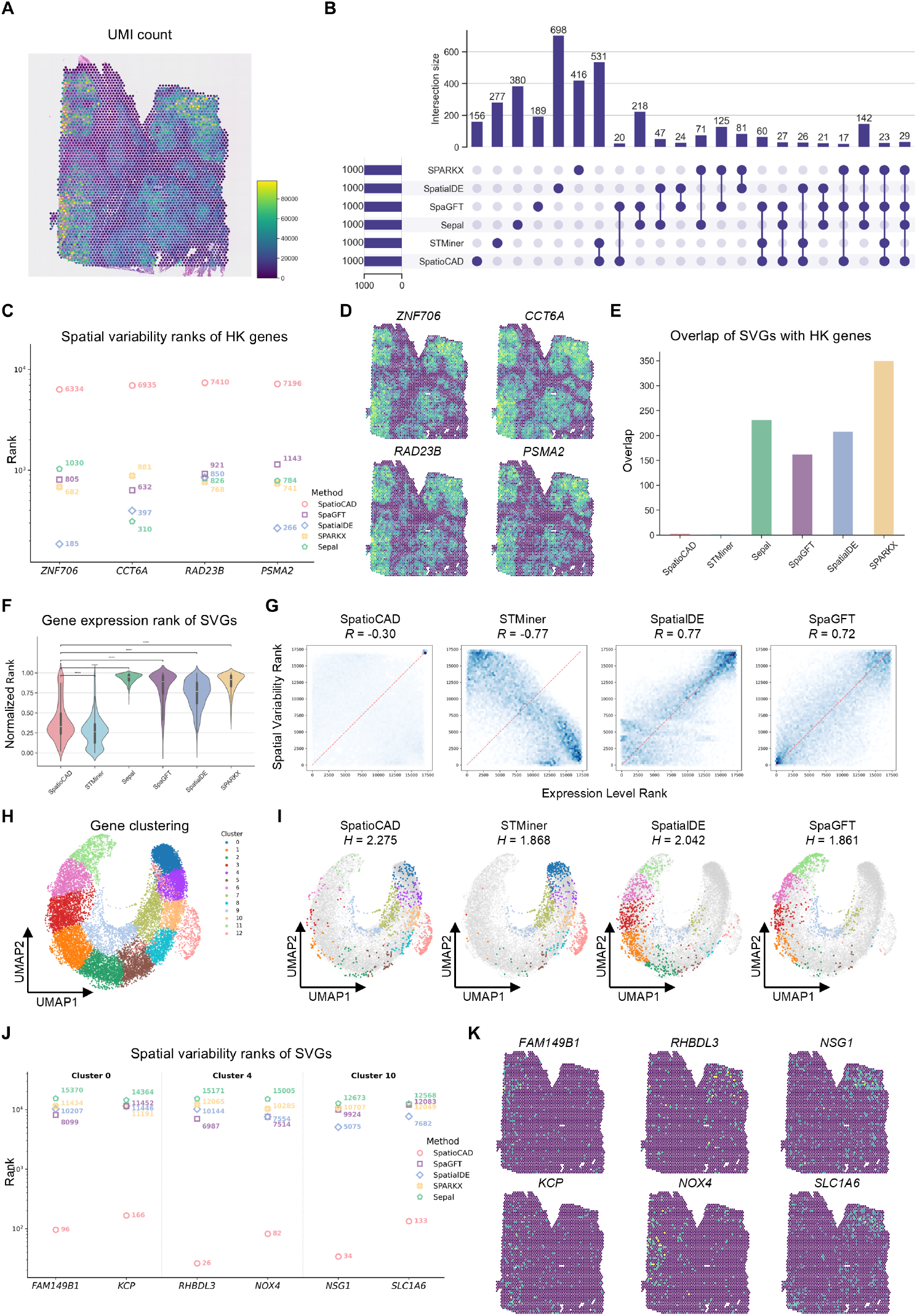
SpatioCAD enables robust identification of functionally diverse SVGs independent of cell density and expression abundance in human lung cancer data. **A** Visualization of the human lung cancer dataset with its UMI counts. **B** UpSet plot shows the overlap among the top 1,000 SVGs identified by the six compared methods (SpatioCAD, STMiner, Sepal, SpaGFT, SpatialDE, SPARKX). **C** Comparison of spatial variability ranks for representative housekeeping genes. **D** Spatial expression patterns of representative housekeeping genes shown in **C. E** The number of overlaps between the top 1,000 SVGs identified by different methods and known housekeeping genes in public databases. **F** Violin plot shows the distribution of normalized expression ranks for the top 1,000 SVGs identified by each method. Wilcoxon rank-sum tests were applied to normalized expression ranks comparing SpatioCAD with other methods (*p*-values = 5.55 × 10^−29^, 2.23 × 10^−266^, 1.29 × 10^−202^, 9.81 × 10^−161^, 1.87 × 10^−239^, respectively). **G** Hexbin density plots show the relationship between gene expression ranks (x-axis) and spatial variability ranks (y-axis). Spearman correlation between the two ranks was calculated for each method (SpatioCAD: 0.30, STMiner: −0.77, SpatialDE: −0.77, SpaGFT: 0.72, SPARK-X: 0.91, Sepal: 0.97). **H** UMAP plot shows the clustering results of all genes based on their expression profiles. **I** Projection of the top 1,000 SVGs identified by each method onto the global gene UMAP. Shannon entropy of their distribution across these clusters was calculated for each method (SpatioCAD: 2.275, STMiner: 1.868, SpatialDE: 2.042, SpaGFT: 1.861, SPARK-X: 1.645, Sepal: 1.203). **J** Comparison of spatial variability ranks for representative genes from clusters 0, 4, and 10. SpatioCAD ranks these genes higher than other methods (SpatialDE, SpaGFT, SPARK-X, Sepal). **K** Spatial expression patterns of representative genes shown in **I**.

Using housekeeping genes as negative controls, we evaluated the specificity of top-ranked SVGs. As shown in the ranking plot (Fig. 6C), several HK genes were spuriously ranked as top SVGs by other methods but were correctly excluded by SpatioCAD by ranking them beyond the top 1000. The spatial expression patterns of these representative HK genes further reveal that their apparent spatial variation closely mirrors the UMI count distribution (Fig. 6D), confirming they reflect cell density variation rather than genuine biological patterning. Quantitatively, SpatioCAD and STMiner yielded the lowest overlap with only 3 and 2 genes detected, respectively. This stands in stark contrast to other methods, which generally exhibited higher overlaps: Sepal (231 genes), SpaGFT (162 genes), SpatialDE (208 genes), and SPARK-X (350 genes) (Fig. 6E). This consistent and robust performance highlights the importance of decoupling cell density to ensure the identification of genuine biological signals in tumors.

Next, we examined whether each method’s top SVGs were systematically biased toward high or low expression by analyzing their normalized expression ranks (**Methods**). SpatioCAD again maintained the lowest bias, with a mean normalized expression rank of 0.396 and a minimal MAD of 0.104 from the ideal value 0.5. In contrast, STMiner favored lowly expressed genes (mean: 0.272; MAD: 0.228), while SpatialDE (mean: 0.735; MAD: 0.235), SpaGFT (mean: 0.826; MAD: 0.326), SPARK-X (mean: 0.883; MAD: 0.383) and Sepal (mean: 0.935; MAD: 0.435) remained heavily skewed towards highly expressed genes (Fig. 6F). Spearman correlation analysis further supported these findings: SpatioCAD showed minimal correlation (*R* = −0.30), effectively decoupling spatial significance from expression abundance, whereas STMiner exhibited a significant negative correlation (*R* = −0.77), and all other methods showed strong positive correlations, ranging from SpaGFT (*R* = 0.72) and SpatialDE (*R* = 0.77) to SPARK-X (*R* = 0.91) and Sepal (*R* = 0.97). Visualization of rank scatter plots confirmed that SpatioCAD’s SVGs were uniformly distributed across expression levels, while SVGs from other methods clustered along diagonal or anti-diagonal patterns, indicating systematic expression-level bias (Fig. 6G, Supplementary Fig. 13).

To assess the functional diversity of identified SVGs, we leveraged the established expression-based clustering framework (**Methods**, Supplementary Figs. 14-18). Genes were clustered based on expression profiles, and Shannon entropy of SVG distribution across the clusters was calculated (Fig. 6H, I). SpatioCAD again achieved the highest entropy (*H* = 2.336), followed by SpatialDE (*H* = 2.032), SpaGFT (*H* = 1.941), STMiner (*H* = 1.833), SPARK-X (*H* = 1.674), and Sepal (*H* = 1.363) (Fig. 6I, Supplementary Fig. 19). This result confirms that SpatioCAD’s capacity to recover a comprehensive spectrum of functional signals is not dataset-dependent but a robust feature of the algorithm.

Analysis of SVG distribution across clusters highlighted method-specific preference (Supplementary Fig. 20). SpatioCAD maintained robust detection of biological signals in all clusters, while SpatialDE, Sepal, SpaGFT, and SPARK-X identified few or no SVGs in low-expression clusters 0, 4, and 10 (Supplementary Fig. 17). In contrast, STMiner exhibited reduced sensitivity in clusters 1, 2, 3, 6, and 11, reinforcing its bias toward lowly expressed genes. These findings corroborate the earlier observed dependency between spatial variability and expression abundance for all the methods.

We further investigated the biological significance of SVGs in the low-expression clusters (Clusters 0, 4, and 10) identified by SpatioCAD but missed by other methods (Fig. 6J). These SVGs revealed two spatially distinct regions within the lung cancer tissue (Fig. 6K). On the left side of the lung cancer tissue, we observed a spatial aggregation of *NOX4, KCP*, and *FAM149B1* (Fig. 6K). This co-localization pattern likely suggests the presence of a tumor-stroma interface undergoing active remodeling. Specifically, *NOX4* has been confirmed to play an important role in pulmonary tissue fibrogenesis [36]. Conversely, *KCP* acts as an enhancer of bone morphogenetic protein (BMP) signaling, a pathway shown to attenuate pulmonary fibrosis [37]. The presence of *KCP* thus suggests that the tissue within this niche exhibits a counter-regulatory response to the fibrotic process. The top-right region, enriched with *NSG1, SLC1A6*, and *RHBDL3*, likely represents the tumor core. Specifically, *NSG1* has been identified as a key gene deeply implicated in the pathology of lung adenocarcinoma, with its expression levels correlating strongly with tumor staging [38]. *SLC1A6* serves as a robust survival predictor, capable of stratifying lung adenocarcinoma patients into high-risk groups with significantly poorer prognosis [39].

#### 2.4.2 SpatioCAD captures more robust biological structures than STMiner

To directly compare methods that address cell density confounding, we conducted a focused analysis between SpatioCAD and STMiner centered on the SVGs uniquely identified by each method. We first quantified the local smoothness of the top SVGs using a local linear regression framework, which predicts the expression level at each spot using its surrounding coordinates (**Methods**). SpatioCAD achieved significantly lower prediction error compared to STMiner (mean error: 5.00 vs 8.26, Wilcoxon-rank test *p*-value = 1.46 × 10^−57^; Fig. 7A). This result indicates that SpatioCAD is capable of filtering out spatially fragmented signals, ensuring the identification of robust and continuous spatial structures.

We next employed Moran’s *I* to assess global spatial autocorrelation (**Methods**). The SVGs identified by SpatioCAD exhibited markedly higher Moran’s I indices than those from STMiner (mean: 0.18 vs 0.07, Wilcoxon-rank test *p*-value = 5.69 × 10^−18^; Fig. 7B). This statistically significant difference confirms that SpatioCAD captures genes with stronger spatial structures, which are more likely to encode rich and interpretable biological information.

We further assessed the statistical stability of identified SVGs using permutation-based testing to evaluate whether their significance persists under random perturbations (**Methods**). SpatioCAD demonstrated exceptional specificity, with all 303 top SVGs exclusive to it achieving statistical significance (*p <* 0.05). In comparison, STMiner yielded a significantly lower number of significant genes (173 genes), with a substantial portion of its top candidates (104 genes) exhibiting high empirical *p*values (*p* ≥ 0.1) (Fig. 7C). This high proportion of non-significant genes suggests that many of STMiner’s candidates may lack genuine spatial variability and represent false positives driven by stochastic noise. Together, these validations on the lung cancer dataset confirm that SpatioCAD generates statistically robust and biologically coherent results, supporting its reliability across diverse cancer types.

**Figure 7.**
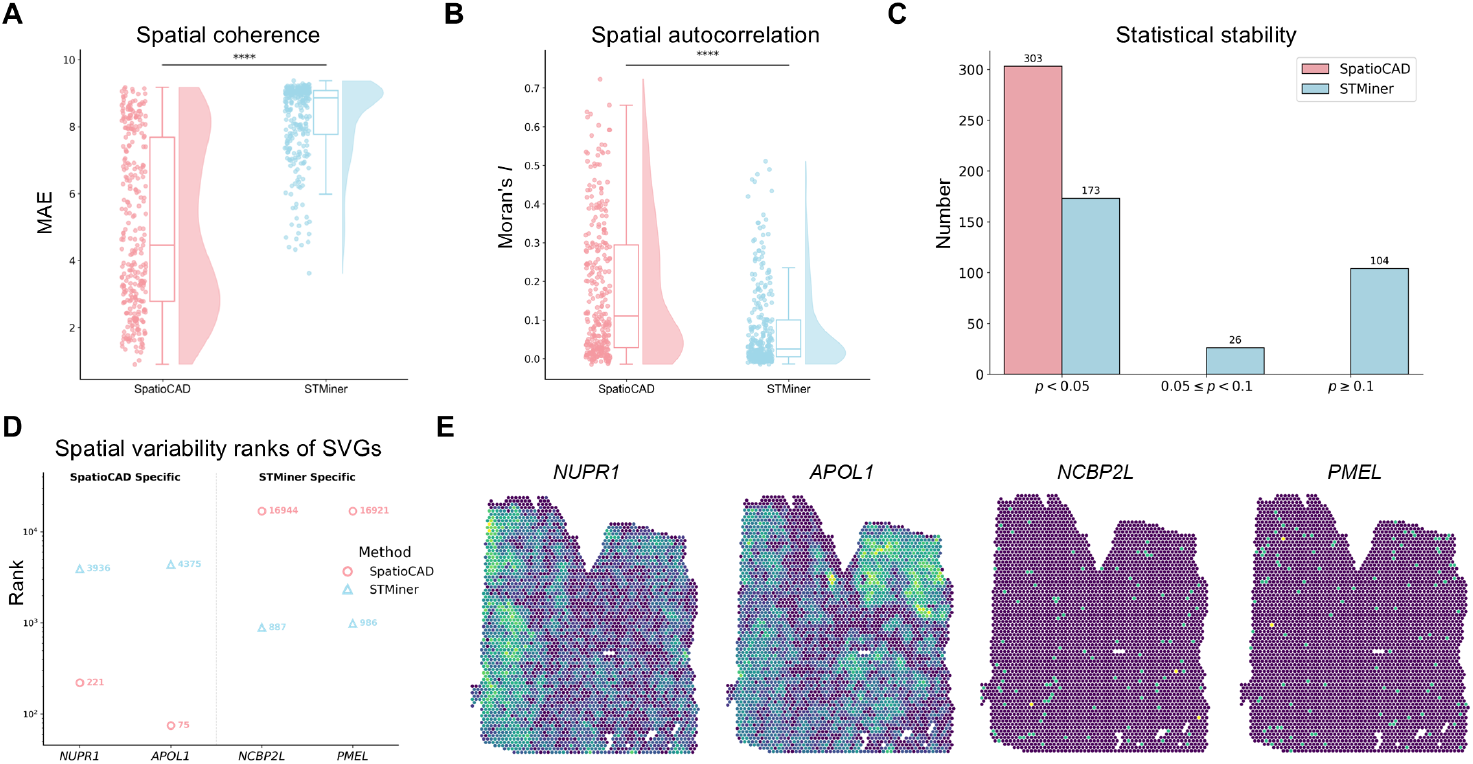
SpatioCAD identifies more spatially coherent and statistically robust SVGs than STMiner in human lung cancer data. **A** Raincloud plot shows the distribution of gene expression reconstruction error for method-specific SVGs. Wilcoxon rank-sum test comparing SpatioCAD with STMiner (*p*-value = 1.46 × 10^−57^). **B** Raincloud plot shows the distribution of global Moran’s *I* indices for method-specific SVGs. Wilcoxon rank-sum test comparing SpatioCAD with STMiner (*p*-value = 5.69 × 10^−18^). **C** Bar plot shows the classification of method-specific SVGs based on permutation-based empirical *p*-values. **D** Comparison of spatial variability ranks for representative method-specific SVGs. **E** Spatial expression patterns of representative genes shown in **D**.

To visually validate these statistical findings, we inspected the expression patterns of several SVGs that were highly ranked by one method but assigned low ranks by another (Fig. 7D). SpatioCAD successfully identified key lung cancer-associated markers, such as *NUPR1* and *APOL1*. Specifically, *NUPR1* has been shown to be essential for tumor progression, as its inhibition significantly suppresses the growth of non-small cell lung cancer [40]. Meanwhile, *APOL1* functions as a critical regulator of autophagy and programmed cell death, influencing tumor cell survival under stress [41]. Visualizing these genes reveals clear, spatially aggregated expression structures (Fig. 7E, left). Conversely, STMiner prioritized some genes characterized by poor spatial prediction and low statistical confidence, such as *PMEL* (empirical *p*-value: 0.40) and *NCBP2L* (empirical *p*-value: 0.40). From a functional perspective, *PMEL* is a pigment cell-specific protein involved in the biogenesis of pigment organelles [42]. Consequently, its identification as a spatially variable gene in lung tissue is highly indicative of a false positive result. Consistent with this functional discrepancy, the spatial patterns for these genes display scattered, discrete noise patterns (Fig. 7E, right), further confirming that SpatioCAD successfully isolates true biological signals from technical artifacts.

Finally, in terms of computational efficiency, SpatioCAD dramatically outperformed STMiner in runtime, completing the analysis in just 40 seconds compared to 23, 410 seconds for STMiner.

### 2.5 SpatioCAD dissects functional heterogeneity in the tumor microenvironment of diffuse midline glioma

One key strength of SpatioCAD is its ability to group SVGs into clusters that share similar spatial expression patterns based on their diffusion profiles, enabling systematic functional dissection of distinct regions within the tumor microenvironment. To demonstrate this capability, we applied SpatioCAD to a publicly available 10X Visium dataset of diffuse midline glioma (DMG). Visualization of three human malignant glioma samples, along with their provided annotations and UMI counts, confirmed the increased cell density in tumor regions (Fig. 8A).

**Figure 8.**
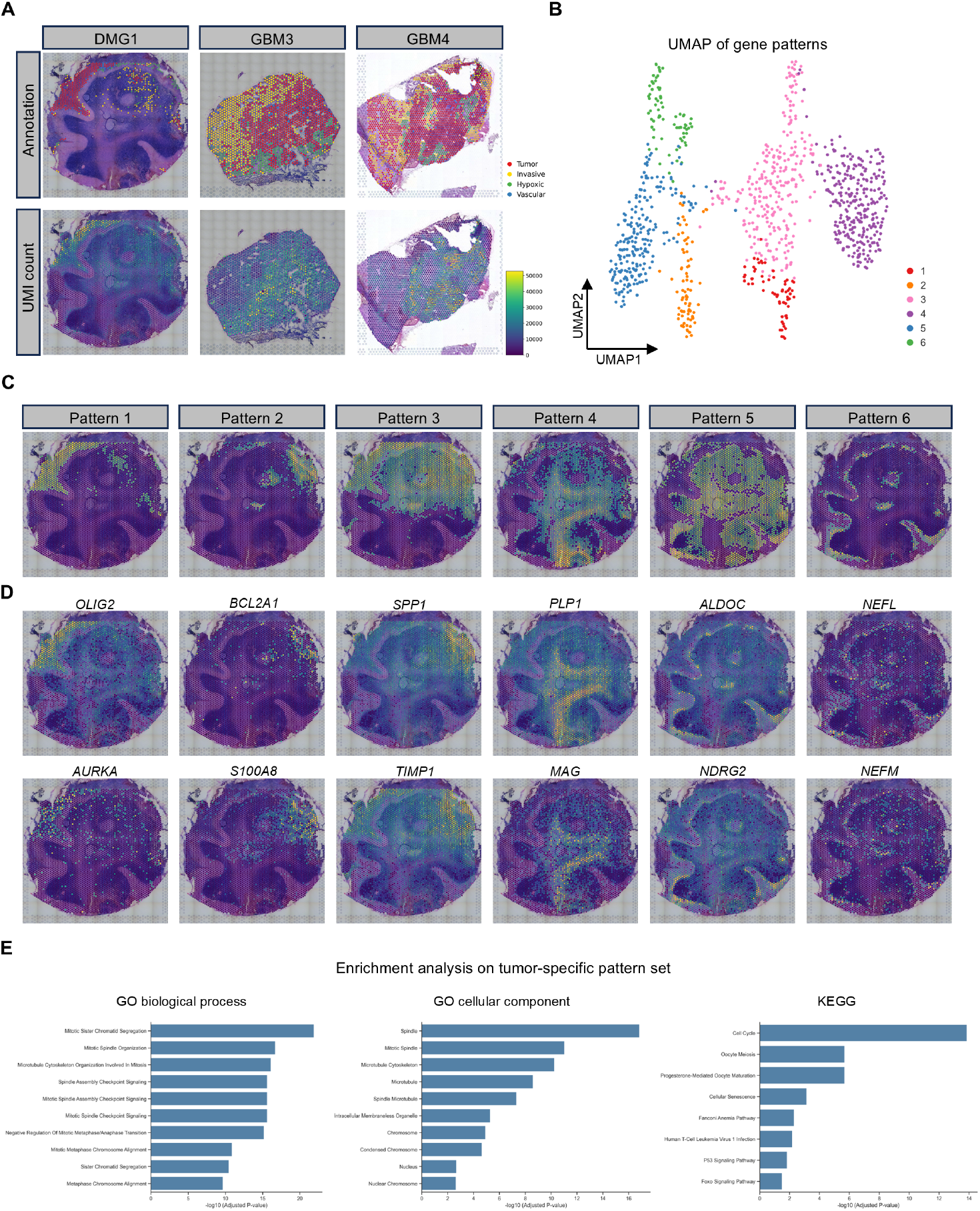
Analysis of human malignant gliomas data. **A** Visualization of three samples from the human malignant gliomas dataset (DMG1, GBM3, and GBM4) with their annotations (upper) and UMI counts (lower). **B** UMAP plot shows the gene clusters based on their spatial distribution characteristics, with genes colored by the cluster labels generated by SpatioCAD. **C** Spatial patterns of the six distinct gene clusters identified by SpatioCAD. **D** Spatial expression patterns of representative genes from each of the six identified clusters. **E** Significantly enriched gene sets (FDR adjusted *p*-value *<* 0.05) in gene set enrichment results for the tumor-specific gene set (Cluster 1) on three reference gene sets: GO biological process, GO cellular component, and KEGG.

We focused on the top 1,000 SVGs identified by SpatioCAD in sample DMG1, which comprises measurements of 36,601 genes across 4,337 spots. These SVGs were clustered based on their spatial diffusion profiles, which model the diffusion process as a superposition of constituent eigenmodes (**Methods**). To evaluate the biological relevance of these spatial diffusion profiles, we visualized the genes in the UMAP embedding constructed from their normalized expression data, colored by SpatioCAD cluster identities (Fig. 8B). The results showed strong concordance: genes belonging to the same SpatioCAD cluster formed compact, distinct groups in expression space, confirming that diffusion profiles effectively capture biologically meaningful spatial patterns. Furthermore, we investigated the spatial patterns of each cluster (Fig. 8C). Comparison with the provided annotations revealed that SpatioCAD successfully captured the tumor core, further validating its power in resolving tissue architecture.

We next analyzed the biological functions of the SVGs within each spatial cluster (Fig. 8D). Pattern 1, which showed high spatial overlap with the annotated tumor core, was enriched for genes integral to the pathogenesis of DMG. These included *OLIG2*, a marker highly expressed in DMG that is essential for the tumor cell propagation [43], and *AURKA*, whose inhibition has been demonstrated to block DMG cell cycle progression [44]. Pattern 2 was predominantly active in the tumor periphery, indicative of a reactive host response to the encroaching malignancy. This cluster was characterized by key inflammatory genes, including *BCL2A1* and *S100A8* [45, 46], a molecular signature characteristic of a robust macrophage response to tumor infiltration [47]. Pattern 3, spanning both the annotated tumor and invasive regions, appeared to represent an active zone of tumor invasion. This observation is supported by previous studies confirming that genes within this cluster, such as *SPP1* and *TIMP1*, are associated with DMG pathogenesis [48, 49].

Beyond characterizing the tumor microenvironment, SpatioCAD also successfully recapitulated the fundamental brain tissue architecture (Fig. 8D). Pattern 4 exhibited a spatial distribution closely corresponding to the inner eosinophilic (pink) regions observed in the H&E-stained histology. This morphological alignment suggested its identity as white matter, which was biologically validated by the specific enrichment of myelin-sheath-associated genes, such as *PLP1* and *MAG* [50]. Pattern 5 spatially coincided with the hematoxylin-rich (dark purple) regions. We thus identified this region as gray matter, a classification strongly supported by the enrichment of astrocyte-specific markers, including *ALDOC* and *NDRG2* [51, 52]. Pattern 6 delineated a distinct, thin layer at the outermost periphery of the tissue. The molecular profile of this pattern was characterized by expression of the neurofilament genes, such as *NEFL* and *NEFM*, indicating the specific capture of a neuronal layer distinct from the inner parenchyma [50, 51].

The gene set enrichment analysis on the tumor-specific cluster (Cluster 1) using the Gene Ontology (GO) and KEGG databases is shown in Fig. 8E. The results indicated an overwhelming enrichment of genes involved in the regulation and machinery of cell division. Specifically, GO biological process analysis showed strong enrichment in mitotic processes, while GO cellular component analysis revealed that the genes were significantly enriched in components of the spindle and mitotic spindle. This finding is highly consistent with uncontrolled proliferation, a core hallmark of cancer [53]. Corroborating this conclusion, the KEGG pathway analysis also identified ‘Cell Cycle’ as the most significantly enriched pathway. We further extended this functional characterization to the remaining clusters (Clusters 2-6), where the enrichment analyses consistently validated our spatial annotations, spanning from inflammatory responses to specific metabolic and synaptic signatures (Supplementary Fig. 21).

Finally, we extended our analysis to the other two malignant glioma samples (GBM3 and GBM4). SpatioCAD consistently resolved the heterogeneous tumor architecture and identified the tumor core in both datasets, confirming the method’s robustness across independent samples (Supplementary Figs. 22-25).

## 3 Discussion

In complex tissues such as tumors, varying degrees of cellular aggregation can lead to marked heterogeneity in cell density. This heterogeneity often complicates the analysis of spatial transcriptomics data, resulting in the misidentification of false-positive SVGs. To address this challenge, we developed SpatioCAD, a two-step computational framework. Leveraging the structural instability of noise signals, SpatioCAD first employs the Roughness Score, derived from standard graph diffusion model, to filter out uninformative genes. Subsequently, it calculates the diffusion time for each gene under the Node-Attributed Graph Diffusion (NAGD) model, a generalization of standard graph diffusion, which effectively decouples spatial patterns from cell density variations, thereby enabling the robust identification of genuine SVGs.

The experimental results demonstrate that SpatioCAD effectively mitigates the false positives caused by cellular heterogeneity while providing unbiased gene selection. This capability facilitates the detection of lowly expressed yet biologically important SVGs, yielding a more diverse and informative SVG set. Furthermore, in a direct comparison with STMiner, an existing method developed for the same purpose, SpatioCAD demonstrates superior efficiency, statistical significance, and robustness. Finally, investigation into the biological functions of the SVGs identified by SpatioCAD reveals its capability to pinpoint clinically significant genes in multiple cancer datasets, confirming its practical relevance.

SpatioCAD is not without limitations. First, unlike many statistical methods, SpatioCAD does not provide a formal statistical measure, such as a *p*-value or *z*-score, for a definitive classification of SVGs and non-SVGs. Second, this framework relies on graph construction and eigen-decomposition of the graph Laplacian matrix. Developing a construction strategy that balances computational efficiency with the preservation of spatial information remains an important direction for future investigation. Lastly, our framework uses the sum of gene expression at each spatial location as a proxy for cell density. While this is a widely adopted and practical strategy, its accuracy can be confounded by technical factors. For example, technical artifacts, such as uneven tissue permeabilization or variations in RNA capture efficiency, can also cause spatial fluctuations in UMI counts that are independent of true cell density changes. Future algorithmic enhancements may benefit from incorporating histology image information to estimate cell density more accurately.

In recent years, identifying cell type-specific SVGs (ct-SVGs) has become an active area of research [54–56]. These methods aim to identify genes that exhibit spatial patterns within specific cell types. While SpatioCAD can be adapted to find variations associated with a particular cell type by using its proportion instead of overall cell density in the NAGD framework, it cannot yet resolve the contributions of multiple cell types simultaneously. Extending SpatioCAD to integrate cell-type deconvolution results represents a promising direction for future research.

## 4 Methods

### 4.1 Context-aware graph diffusion model for pinpointing spatially variable genes in heterogeneous tissues

SpatioCAD is a computational framework designed to identify spatially variable genes (SVGs) within heterogeneous tissues, such as tumors. The core principle of SpatioCAD is to model the spatial diffusion process of gene expression in complex tissues, based on the assumption that genes with structured spatial patterns will reach a steady state more slowly than those with random distributions. The framework consists of two main stages. First, a Roughness Score (RS), derived from the standard graph diffusion model, is calculated to filter out genes with noisy expression patterns, thereby preventing them from confounding downstream analyses. Second, a node-attributed graph diffusion model is proposed to simulate signal propagation across the heterogeneous tumor microenvironment. The spatial variability of each gene is then quantified by the diffusion time required to reach a steady state, with genes exhibiting stronger spatial structure showing longer diffusion durations. Theoretical proofs for this framework are provided in the Supplementary Note.

#### 4.1.1 Spatial graph construction

Let **X** ∈ ℛ^*N* ×*G*^ be the preprocessed spatial transcriptomics data matrix, where *N* is the number of spatial locations and *G* is the number of genes. The spatial coordinates of the locations are denoted by **s** ∈ ℛ^*N* ×*D*^, with the dimensionality *D* = 2 or 3. First, SpatioCAD constructs an undirected graph **G** = (*V, E*), where the nodes *V* represent the *N* locations and the edges *E* are established using the mutual *k*-nearest neighbor principle. Specifically, an edge *e*_*ij*_ exists between nodes *i* and *j* if and only if *i* ∈ 𝒩_*k*_(*j*) and *j* ∈ 𝒩_*k*_(*i*), where 𝒩_*k*_(*j*) denotes the set of *k*-nearest neighbors of node *j*. A symmetric adjacency matrix **A** = (*a*_*ij*_) ∈ ℛ^*N* ×*N*^ is then constructed, where the edge weights are calculated using a self-tuning adaptive Gaussian kernel [57]:

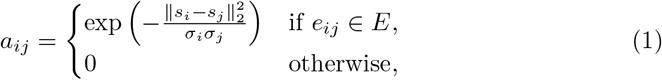

where ∥ · ∥_2_ denotes the Euclidean distance. The local scale parameter *σ*_*i*_ adapts to the local neighborhood density and is defined as the Euclidean distance from location *i* to its *k*^′^-th nearest neighbor.

Based on simulation experiments (Supplementary Fig. 26), the default values of *k* and *k*^′^ were set to 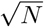 and 0.5 log *N*, respectively, to balance performance and computational efficiency.

#### 4.1.2 Noise gene filtering

To enhance the statistical power and biological relevance of SVG identification, we implement a filtering strategy based on the standard graph diffusion model to exclude genes dominated by noise. Here, each gene’s expression profile is modeled as a graph signal on a constructed spatial graph, where nodes correspond to distinct spatial locations. We denote the expression level of any gene as **x** ∈ ℛ^*N* ×1^, with *x*_*i*_ being the expression value at spot *i*. The diffusion process of these signals is governed by the graph heat equation [58, 59]:

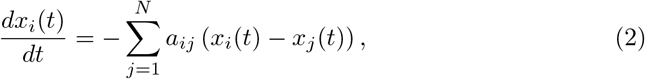

where *x*_*i*_(*t*) denotes the signal state of location *i* at time *t*, with the initial state **x**(0) representing the observed gene expression in **X**. We can express this in vector form:

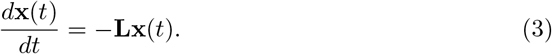

Here, **L** is the graph Laplacian matrix defined as **L** = **D**−**A**, where **A** is the previously defined adjacency matrix and **D** is the diagonal degree matrix with entries *d*_*ii*_ = ∑_*j*_ *a*_*ij*_. This differential equation has a well-defined analytical solution:

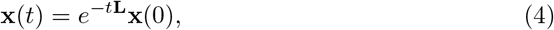

which precisely describes the state of the gene signal at any given time *t*.

Biologically meaningful patterns are typically characterized by local consistency, while noise signals tend to change drastically during the initial diffusion steps. Thus, we define a Roughness Score (RS) to filter the noise genes. This score is formally derived from the *L*_2_-norm of the initial signal variation:

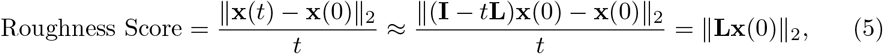

where **x**(0) and **x**(*t*) denote the initial and time-*t* state signal vectors, respectively. The approximation is obtained using a first-order Taylor expansion of Equation (4). In practice, we may directly set *t* = 1 and use ∥**x**(*t*) − **x**(0) ∥_2_ or 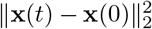 as the RS. Notably, from the perspective of spectral graph theory, a higher RS indicates that the signal is dominated by high-frequency components on the graph (Supplementary Note).

To establish an adaptive filtering threshold, we model the distribution of the Roughness Scores across all genes using a Gaussian Mixture Model (GMM). We fit GMMs with varying component numbers (by default, from 1 to 10) and select the optimal model based on the Bayesian Information Criterion (BIC) [60]. The Gaussian component with the highest mean RS is then identified as the noise component. Finally, we calculate the posterior probability of belonging to this noise component for each gene. Genes with a posterior probability exceeding a pre-defined threshold (by default, 0.9) are classified as lacking sufficient spatial information and are subsequently excluded from further analysis.

#### 4.1.3 Gene spatial variability quantification

Complex biological tissues, such as tumor tissues, exhibit significant heterogeneity in cellular composition. However, standard graph diffusion models assume uniform node properties, limiting their applicability in heterogeneous contexts. Thus, we propose a Node-Attributed Graph Diffusion (NAGD) model that explicitly incorporates node-specific attributes. The NAGD model extends the conventional diffusion framework by integrating the following key biological principles. First, consistent with canonical models, diffusion between nodes is affected by their spatial connectivity, as encoded in the adjacency matrix **A**. Second, to account for tissue heterogeneity, we assume the diffusion process is driven by the difference in signal concentration (*x*_*i*_*/n*_*i*_ for node *i*) rather than raw signal intensity. This concentration difference between paired nodes is represented by (*x*_*i*_*/n*_*i*_ − *x*_*j*_*/n*_*j*_). Finally, the diffusion process is further constrained by the number of potential communication channels between nodes, denoted as *w*_*ij*_, which is related to the cellular populations of the two interacting nodes.

Integrating these principles yields the following equation:

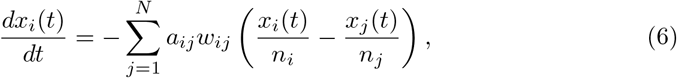

where *n*_*i*_ denotes the cell density of node *i*, which we approximate as the sum of normalized expression across all genes at node *i*. A reliable choice for *w*_*ij*_ is the arithmetic mean of cell densities, (*n*_*i*_ + *n*_*j*_)*/*2. Notably, our model represents a generalization of the standard graph diffusion model, to which it reduces when cell numbers across all nodes are uniform.

By setting 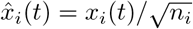, Equation (6) can be rewritten as:

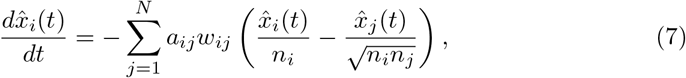

which can be expressed in vector form as:

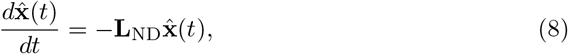

with **L**_ND_ defined as:

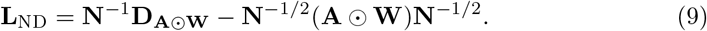

In this expression, ⊙ denotes the Hadamard (element-wise) product, **D**_**A**⊙**W**_ is the degree matrix derived from the element-wise product of **A** and **W**, and **N** = diag(*n*_1_, *n*_2_, …, *n*_*N*_) is the diagonal matrix of the cell density vector. Analogous to Equation (4), this linear equation has the analytical solution:

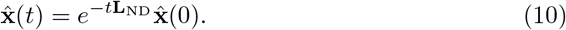

To further simplify the analytical solution, we perform a spectral decomposition of the NAGD Laplacian matrix **L**_ND_:

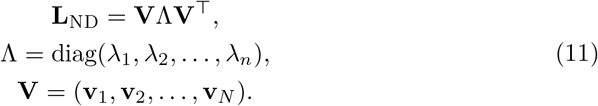

Here, for a connected graph, 0 = *λ*_1_ *< λ*_2_ ≤ · · · ≤ *λ*_*N*_ are the eigenvalues of **L**_ND_, and **V** is the orthogonal matrix whose columns **v**_*k*_ are the corresponding eigenvectors, with the first eigenvector **v**_1_ corresponding to the steady state (Supplementary Note 2).

By substituting this spectral decomposition into Equation (10), the diffusion process can be expressed as a sum of its constituent eigenmodes:

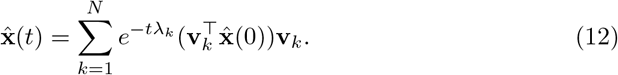

As *t* → ∞, the system converges to its steady state, which is determined by the eigenvector corresponding to the single zero eigenvalue (*λ*_1_ = 0):

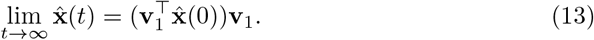

Provided all genes are first normalized to uniform total expression, their diffusion processes will converge to an identical steady state, underpinned by the following two properties. First, for a symmetric matrix **W** = (*w*_*ij*_), the total signal sum, 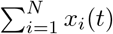, is conserved throughout the diffusion process (Supplementary Note 2). Second, the system evolves to a unique steady state **x**^∗^, which is directly proportional to the cell density vector: **x**^∗^ = *c* · **n**, where **n** = (*n*_1_, *n*_2_, …, *n*_*N*_)^⊤^ and *c* is a scalar constant (Supplementary Note 2). This provides a common endpoint, making their rates of convergence directly comparable. We therefore introduce a characteristic diffusion time as a robust metric to quantify the spatial variability of each gene.

We formally define the characteristic diffusion time, *t*^∗^, as the time required for the change rate norm of the diffusion state vector, 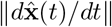, to fall below a predefined threshold *ϵ*:

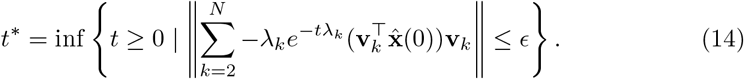

Given the orthogonality of the eigenvectors, this expression simplifies under the *L*_2_norm to:

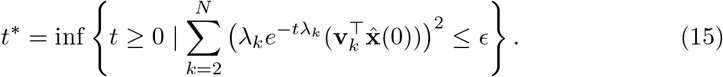

In practice, the convergence rate is governed by the slowest-decaying terms, which correspond to the smallest non-zero eigenvalues. To obtain a tractable analytical expression, we approximate *t*^∗^ by considering the *m* (by default, *m* = 50) slowest-decaying eigenmodes (from *λ*_2_ to *λ*_*m*+1_), which dominate the diffusion process. This simplification yields the final closed-form approximation:

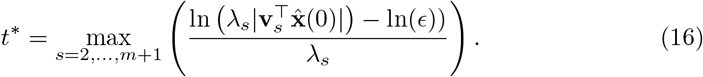

The characteristic diffusion time thus provides a quantitative estimate of spatial variability for each gene, where a larger *t*^∗^ value implies a more significant spatial pattern.

To enhance sensitivity to spatial patterns across the wide dynamic range of gene expression, we introduce a rank aggregation strategy. We apply a series of *p*-th power transformations (by default, *p* ∈ {1, 1.5, 2}) to the normalized expression matrix before computing the diffusion time, thereby generating a distinct gene spatial variability ranking for each transformed version. These rankings are then integrated using the lexicographical ordering method to produce a final, robust rank of spatial variability for all genes.

#### 4.1.4 Computational complexity

The computational complexity of SpatioCAD is primarily determined by four key steps. First, constructing the spatial graph is dominated by the *k*-nearest neighbor search, with a complexity of *O*(*N* log *N*). Second, calculating the Roughness Score for all *G* genes involves a single sparse-dense matrix multiplication, with a complexity of *O*(*kGN*). Third, a key computational advantage of SpatioCAD is that it only requires the *m* eigenmodes corresponding to the smallest non-zero eigenvalues of **L**_ND_. These are computed efficiently using iterative solvers with an approximate complexity of *O*(*kmN*), thus avoiding the costly *O*(*N* ^3^) required for a full decomposition. Finally, computing the diffusion time *t*^∗^ for all *G* genes, as defined in Equation (16), has a complexity of *O*(*mGN*). Therefore, the overall computational complexity of the SpatioCAD pipeline is approximately *O*((*k* + *m*)*GN*), demonstrating that the method scales linearly with the number of genes (*G*) and spatial locations (*N*).

### 4.2 Compared methods

To evaluate the performance of SpatioCAD, we benchmarked it against five existing methods for identifying SVGs: STMiner [19], Sepal [15], SpaGFT [14], SpatialDE [4] and SPARK-X [13]. These methods were selected to represent a diverse range of underlying algorithms and statistical models.

To ensure a fair comparison, we filtered out spots with ≤ 1 detected genes and genes expressed in ≤ 100 spots. Each method was subsequently run on the common filtered expression matrix using its default parameters.

**STMiner** [19] is a method that leverages optimal transport theory to identify SVGs. This method is available within the Python package `STMiner`. We applied the core function `spatial_high_variable_genes` to log-transformed expression counts. Genes were ranked in descending order based on the resulting z-score.

**Sepal** [15] models spatial expression patterns using Fick’s second law of diffusion. This method is available within the Python package `sepal`. The analysis was performed with the function `models.propagate` on normalized and scaled data. Genes were ranked in descending order based on the resulting time.

**SpaGFT** [14] utilizes a graph Fourier transform to identify SVGs based on the Fourier mode. This method is available within the Python package `SpaGFT`. Following standard library size normalization and log-transformation of the count data, we ran the function `detect_svg` to obtain scores. Genes were ranked in descending order based on the resulting `gft_score`.

**SpatialDE** [4] employs Gaussian processes to model and identify genes with significant spatial variation. This method is available within the Python package `SpatialDE`. Following the standard protocol, we first stabilized the data variance (NaiveDE.stabilize) and regressed out the library size effect (`NaiveDE.regress_out`) before running the main function SpatialDE.run. Genes were ranked in ascending order based on the resulting qval.

**SPARK-X** [13] is a non-parametric method that uses a robust kernel-based statistical test to detect diverse spatial patterns. This method is available within the R package `SPARK`. We performed the analysis using the function `sparkx` with the option `mixture`. Genes were ranked in ascending order based on the resulting adjustedPval.

### 4.3 Datasets

#### 4.3.1 Simulation datasets

To simulate the tumor microenvironment, we utilized the reference-free module of SRTsim [61] to generate a square-shaped tissue slice containing 1079 randomly distributed spots. To evaluate algorithmic robustness, we implemented two different scenarios: one with square-shaped domains and another with strip-shaped domains (Fig. 2A). The tissue slice was spatially partitioned into five distinct domains, denoted as A, B, C, D, and E. In both scenarios, domain E was designated as the tumor core, while domains A-D represented the normal tissue.

We then simulated the gene expression counts by sampling from a Zero-Inflated Negative Binomial (ZINB) distribution using SRTsim, with global parameters set to a zero percentage of 0.01 and a dispersion of 0.3. These genes were assigned to three functional groups: 900 background genes, 200 SVGs, and 50 noise genes.

To disentangle the contributions of heterogeneity in cell density from the diverse spatial expression patterns of genes, we defined the fold-change (FC) for each gene *g* at a given spot *i* as a product of two components: cell density, *ρ*(*i*), and Spatial Variability, SV_*g*_(*i*). The relationship is expressed as:

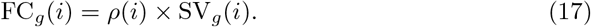

Here, the cell density *ρ*(*i*) is solely dependent on the spatial location and is designed to simulate variations in cell density, such as the hypercellularity in tumor tissues. We therefore set *ρ*(*i*) to 10 for spots in domain E (tumor) and 1 for spots in domains A-D (normal). The second component, SV_*g*_(*i*), is a gene-specific parameter, which governs the intrinsic spatial expression pattern of each gene. Accordingly, we assigned specific SV_*g*_(*i*) values for genes in the three predefined groups. For each scenario, we simulated data under two conditions: with and without noise genes.

##### Background Genes

For background genes, the SV value was maintained at a constant of 1 across all domains. This configuration results in a final FC vector of [1, 1, 1, 1, 10] across domains [A, B, C, D, E], thereby simulating an increase in transcript abundance driven solely by the higher cell density in the tumor domain. To model varying expression levels, these genes were assigned different base mean expression values, stratifying them into a high-expression group (base mean of 4) and a low-expression group (base mean of 2).

##### SVGs

The SVGs were assigned a base mean expression of 2 and stratified into two categories: high-signal (100 genes) and low-signal (100 genes). High-signal SVGs were characterized by higher SV ratios in the designated domains (3.0 for domain B, 2.0 for domain C, 2.5 for domain D, and 2.0 for domain E). Conversely, low-signal SVGs exhibited lower SV ratios in each designated domain with the same predefined domain-specific SV ratios. The resulting FC vectors for these SVGs, which reflect the complex interplay between cell density and intrinsic gene activity, are illustrated in Fig. 2B.

##### Noise Genes

To simulate non-informative signals, we employed the noise gene generation module in SRTsim, setting a low base mean expression of 0.1. This module distributes expression counts randomly across all spots, independently of the predefined domain architecture and cell density variations. These genes therefore serve as a negative control, representing random biological or technical noise.

#### 4.3.2 Real datasets

We applied SpatioCAD and the five compared methods to three publicly available spatial transcriptomics datasets.

##### Human breast cancer data

This dataset, generated from human breast cancer tissue using the 10X Visium platform, is publicly accessible through the 10X Genomics public data repository [23] (https://www.10xgenomics.com/products/xenium-in-situ/preview-dataset-human-breast). It initially comprised 18,085 genes profiled at 4,992 spatial locations. We retained all locations with non-zero expression for at least one gene and filtered out genes expressed in fewer than 100 locations, yielding a final set of 17,633 genes across 4,992 locations for analysis.

The dataset is accompanied by detailed annotations provided by 10X Genomics. In our analysis, we focused on all 11 annotated cell types: invasive, mixed/invasive, DCIS #1, DCIS #2, stromal, immune, adipocytes, stromal/endothelial, mixed, myoepithelial/stromal/immune, and stromal/endothelial/immune.

##### Human lung cancer data

This dataset was generated from human lung cancer tissue using the 10X Visium platform, and is publicly available from the 10X genomics public datasets (https://www.10xgenomics.com/datasets/human-lung-cancer-ffpe-2-standard). It initially comprised 18,085 genes measured across 3,858 spatial locations. With the same filtering method, we retained a final set of 17,269 genes across 3,858 locations for analysis.

##### Human malignant gliomas data

It was generated from human malignant glioma tissue using the 10X Visium platform, and is publicly available from the Gene Expression Omnibus (GEO: GSE194329) [62], which contains five diffuse midline glioma (DMG) and five glioblastoma (GBM) samples. Our analysis focused on the DMG1 sample (sample id: DMG1_short read spatial transcriptomics), with the raw data containing 36,601 genes measured across 4,337 spatial locations. We retained a final set of 15,448 genes across 4,337 locations for analysis with the same filtering method as above. The original study provides annotations for four distinct histopathological regions within the tumor microenvironment: Tumor, Invasive, Hypoxic, and Vascular.

### 4.4 Details of analyses

#### 4.4.1 Data preprocessing

We performed quality control (QC) on the raw count matrix using the Scanpy package [63]. We removed genes expressed in fewer than 50 (simulated datasets) or 100 (real datasets) locations, as well as locations with no gene detection. This filtered matrix then served as the common input for all methods.

Following this common QC, subsequent preprocessing steps diverged according to the recommended guidelines for each compared method. For SpatioCAD, each gene’s expression vector was normalized such that its total counts summed to 10,000 across all locations to preserve the underlying biological variability. This normalized expression matrix was log-transformed before being input into the model.

#### 4.4.2 Evaluation on simulation data

To quantitatively assess and compare the performance of each method in identifying SVGs, we focused on two key metrics: statistical power (Power) and the False Discovery Rate (FDR). Power (sensitivity) was defined as the fraction of true SVGs correctly identified by a method, while FDR represented the proportion of non-SVGs among all genes declared significant. We generated the evaluation curves by calculating the power at five predefined FDR thresholds: 0.01, 0.03, 0.05, 0.075 and 0.1. A superior method is expected to achieve higher power for any given FDR threshold.

#### 4.4.3 Evaluation on real data

##### Overlap between SVGs and housekeeping genes

To evaluate the propensity of each method to identify false positives, we measured the overlap between the top 1,000 identified SVGs and the HRT Atlas v1.0 database of human housekeeping genes [25]. A higher overlap was considered indicative of a greater likelihood of false positives, as housekeeping genes are generally not expected to exhibit spatial variability.

##### Expression-dependent bias in SVG detection

To assess potential expression-dependent bias, we performed a two-stage evaluation of the relationship between spatial variability and gene expression levels. First, we examined the distribution of expression ranks of the top 1,000 SVGs identified by each method relative to all filtered genes. Specifically, all filtered genes were sorted in descending order based on their mean expression level. The rank of each gene *g*, Rank_*g*,original_, was subsequently normalized to a range of [0, 1]:

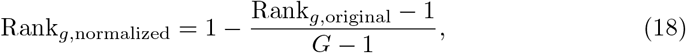

where *G* represents the total number of filtered genes and Rank_*g*,normalized_ thereby quantifies a gene’s relative expression standing. In this framework, a normalized rank of 0 corresponds to the lowest expression value, while 1 indicates the highest. An unbiased method is expected to yield SVGs with a mean normalized rank of approximately 0.5.

Second, to quantify the dependence of SVG detection on expression level, we calculated the Spearman correlation coefficient [64] (*R*) between the rank of spatial variability and mean expression level of the genes:

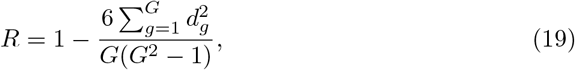

where *d*_*g*_ represents the difference between the two ranks of gene *g*, defined as *d*_*g*_ = Rank_*g*,original_ − Rank_*g*,variability_. A correlation coefficient *R* close to 0 indicates that the method’s prioritization of SVGs is independent of their abundance. A positive correlation indicates that genes with higher expression levels tend to exhibit greater spatial variability, and vice versa.

##### Functional diversity of SVGs

To verify whether the identified SVGs capture a broad spectrum of biological functions, we quantified their functional diversity based on gene clustering. Specifically, all genes were first embedded into a lowdimensional space using Principal Component Analysis (PCA) and Uniform Manifold Approximation and Projection (UMAP) [65]. The Leiden algorithm [66] was then applied to cluster the genes into *M* distinct co-expression modules, denoted as 𝒞 = {*C*_1_, *C*_2_, …, *C*_*M*_} . We assessed the biological relevance of each cluster through gene set enrichment analysis using the Gene Ontology (GO) database, confirming that these modules correspond to distinct biological functions.

A method capable of capturing functionally diverse SVGs should distribute them across multiple independent modules, rather than concentrating them toward a narrow functional subset. For each method, we calculated the probability distribution of the SVGs across these clusters. Functional diversity was then quantified using the Shannon entropy [67] (*H*):

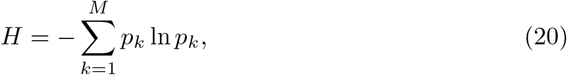

where *p*_*k*_ is the proportion of SVGs belonging to cluster *C*_*k*_. A higher entropy value indicates that the identified SVGs are more evenly distributed across different functional clusters, reflecting the method’s ability to recover diverse spatial-functional patterns.

##### Spatial coherence of SVGs

To further assess the quality of SVGs, we evaluated the spatial coherence of the identified SVGs using a neighborhood-based prediction model. This framework is based on the assumption that biologically genuine spatial patterns should exhibit sufficient local smoothness, such that the expression level at any given location can be reliably predicted from its immediate spatial neighbors.

The evaluation was executed in two steps for each location *i*. First, a local linear regression model was trained using exclusively its neighbors, 𝒩_*k*_(*i*), to capture the local relationship between spatial location and gene expression. Second, we used this fitted model to predict the gene expression at the central location of spot *i* itself. The corresponding prediction error quantifies the local coherence of the expression pattern.

Specifically, for a given gene expression *x*_*i*_ at location *i*, the local linear parameter estimates, (*β*_0_, *β*_1_), are obtained by minimizing the following least squares objective function:

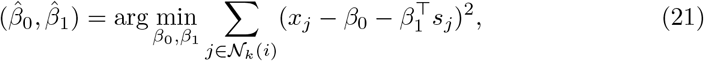

where 𝒩_*k*_(*i*) is the set of *k* = 15 nearest neighbors of location *i*, and *s*_*j*_ is the spatial coordinate vector of location *j*. The estimated parameters, 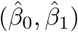, capturing the local expression trend, were then used to predict the expression at the target location *i*:

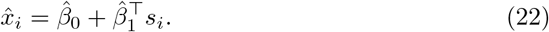

The overall coherence of a given gene and its local neighborhood regions was quantified using the Mean Absolute Error (MAE) of the prediction across all *N* locations:

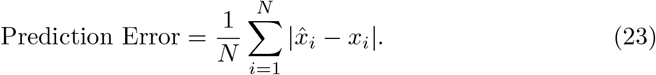

Here, a lower MAE signifies superior coherence, indicating that the gene’s spatial pattern can be reconstructed from its local neighborhood structure.

##### Spatial autocorrelation of SVGs

To validate the spatial significance of the detected genes from a global perspective, we calculated Moran’s *I* statistic [32] for the top-ranked SVGs. This metric serves as a standard measure of spatial autocorrelation, quantifying the extent to which gene expression exhibits clustered spatial patterns rather than a random distribution. A higher average Moran’s *I* indicates a superior capability to prioritize genes with stable spatial structures.

##### Statistical stability of SVG detection

To assess the stability of the detected SVGs for each method, we employed a permutation-based testing framework. For a given SVG identified by a specific method, we randomly shuffled its spatial coordinates while holding the expression profiles of all other genes fixed. The rank of spatial variability in descending order for the permuted gene was then calculated. We quantified statistical stability using an empirical *p*-value derived from *N*_perm_ permutations (by default, *N*_perm_ = 50):

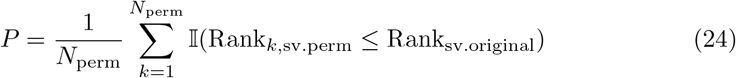

where Rank_sv.original_ denotes the spatial variability ranking of the gene in the observed data, Rank_*k*,sv.perm_ represents its rank in the *k*-th permutation, and 𝕀(·) is the indicator function. A lower *p*-value indicates higher statistical stability of the identified spatial pattern.

##### SVG clustering

To systematically categorize the identified SVGs into distinct spatial modules, SpatioCAD employs a clustering step leveraging their spatial diffusion profiles. The spectral decomposition of the diffusion process (Equation (12)) provides a natural feature space for gene representation. Specifically, the projection of a gene’s expression 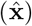 onto the *k*-th eigenvector (**v**_*k*_), quantified by the coefficient 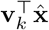, measures the alignment of the gene with the corresponding eigenmode. To group genes with similar spatial topologies, we constructed a low-dimensional spectral embedding, 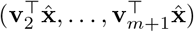, and subsequently applied *k*-means clustering to these feature vectors.

Following the identification of distinct gene modules, SpatioCAD derived a representative spatial pattern for each gene set. First, we computed an expression ratio map by calculating the fraction of genes within the cluster that are expressed at each location. The final spatial pattern was then generated by applying a threshold to this map: specifically, the value at any given location was set to the cluster’s mean expression if its local expression ratio exceeds a defined cutoff (by default, 20%); otherwise, it was assigned a value of zero.

##### UMAP visualization

For the human malignant gliomas dataset, we projected the normalized spatial expression profiles of the top 1,000 SVGs into a two-dimensional space using the Uniform Manifold Approximation and Projection (UMAP) [65]. This visualization aimed to validate the coherence of our previously identified gene clusters. In the resulting UMAP embedding, each point corresponds to a gene, and the proximity between points is indicative of the similarity in their spatial expression profiles.

##### Gene set enrichment analysis

We performed gene set enrichment analysis using the Python package `gseapy` [68], querying against several gene set libraries: `GO_Biological_Process_2025, GO_Cellular_Component_2025`, and `KEGG_2021_Human`. Enriched terms with an adjusted *p*-value *<* 0.05 were considered statistically significant.

## 4.5 Code availability

The SpatioCAD software code is publicly available at https://github.com/kotone-429/SpatioCAD.

## 5.6 Data availability

The data analyzed in this study can be accessed through: Human breast cancer data: https://www.10xgenomics.com/products/xenium-in-situ/preview-dataset-human-breast; Human lung cancer data: https://www.10xgenomics.com/datasets/human-lung-cancer-ffpe-2-standard; Human malignant gliomas data: Gene Expression Omnibus (GEO) database with accession code `GSE194329`.

